# Dissecting Medullary Raphe Neurons Regulating Multiple Thermogenic Pathways

**DOI:** 10.1101/2025.09.01.673572

**Authors:** Shuntaro Uchida, Mitsue Hagihara, Kenichi Inoue, Takaya Abe, Takeshi Sakurai, Kazunari Miyamichi

## Abstract

Thermogenesis is critical for survival and health in mammals. Although the thermoregulatory systems in the preoptic area are well documented, the downstream processing of these central signals—particularly by medullary neurons involved in the control of shivering and sympathetic activation of brown adipose tissue (BAT)—remains incompletely understood. Here we show that *vesicular glutamate transporter type 3* (*vGluT3*)-expressing neurons in the medullary raphe pallidus (RPa) become active immediately before a spontaneous increase in body temperature. These neurons remain inactive under experimentally induced hypometabolic conditions and are necessary for rapid recovery from hypothermia. Furthermore, they communicate with multiple brainstem systems involved in the integration of thermal cues. Notably, RPa-*vGluT3* neurons can drive shivering via specific brainstem premotor neurons, in addition to regulating sympathetic outflows for BAT thermogenesis and heat-conserving piloerection. These data indicate that RPa-*vGluT3* neurons function as medullary hubs, coordinating sympathetic and somatic motor outputs to increase body temperature.

**Graphical abstract:** 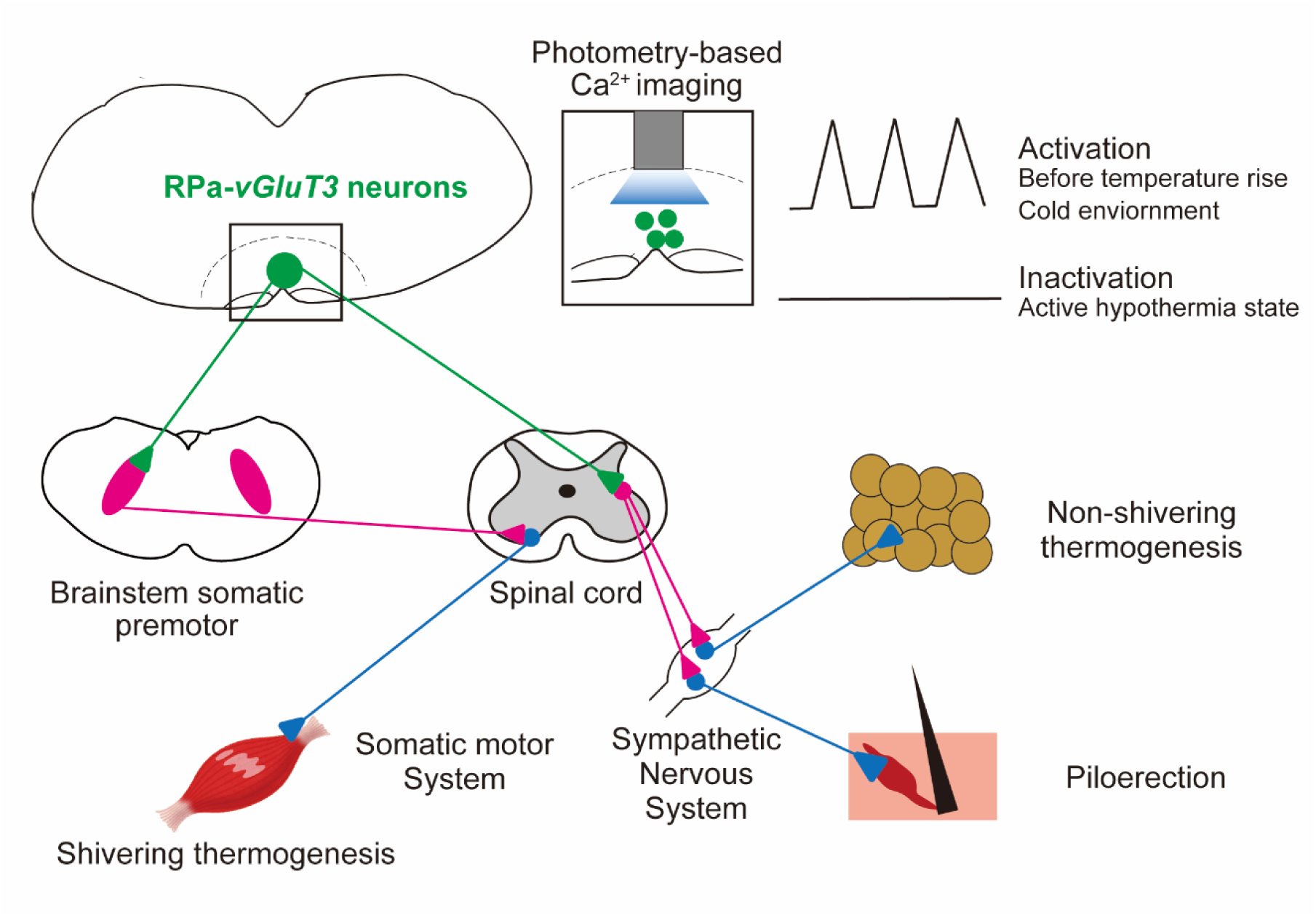

**Highlights:**

- Raphe pallidus-*vGluT3* neurons are active immediately preceding a spontaneous rise in body temperature.
- They are suppressed during the state of Q-neuron–induced hypothermia and hypometabolism (QIH).
- They connect with multiple brainstem pathways involved in thermoregulation.
- They control shivering via brainstem somatic premotor neurons.

## Introduction

Mammals possess a remarkable capacity to tightly regulate their core body temperature (T_c_) within a specific range, typically maintaining it above that of the surrounding environment. The preoptic area (POA) is instrumental in the thermal regulation process. It receives and integrates diverse inputs from various thermosensory systems located in the skin, abdominal organs, and spinal cord, and directly measures local brain temperature^1–3^. When POA neurons detect deviations in T_c_ from their set point or anticipate thermal changes based on environmental cues, they adjust thermogenesis by activating or inactivating brown adipose tissue (BAT), skeletal muscle shivering, and vasoconstriction^1, 2^. These regulatory responses are primarily governed by neural pathways connecting the POA to the medullary regions that control autonomic and/or skeletal thermogenic functions^2^. Over the past decade, studies in rodent models have revealed the molecular identities of central thermoregulatory neurons in the POA and mapped their input/output neural circuit organizations. However, our understanding of medullary thermoregulatory systems—particularly their cellular organization, activity dynamics, and neural networks—remains limited.

Specifically, recent advancements in viral genetic tools in mice^4^ have greatly enhanced our knowledge of central thermoregulatory neurons in the POA^1, 5^. For instance, specific subsets of POA neurons, such as those expressing the prostaglandin E type 3 receptor (EP3R), are implicated in fever responses triggered by lipopolysaccharides or endogenous pyrogenic mediators such as prostaglandin E2 (PGE2)^6–10^. In this context, immune signals triggered by lipopolysaccharides are critically mediated through a defined inhibitory neural population located in the ventral medial preoptic area^11^. POA neurons expressing the transient receptor potential cation channel M-type 2 (TRPM2) may play a role in detecting local POA temperature^12^. Warmth-sensing POA neurons coexpressing pituitary adenylate cyclase-activating polypeptide (PACAP) and brain-derived neurotrophic factor (BDNF) can induce hypothermia upon activation^13^. Moreover, POA neurons expressing the leptin receptor (LeptR)^14^, violet light-sensitive opsin 5 (OPN5)^15^, pyroglutamylated RFamide peptide (QRFP)^16^, and those active during daily torpor^17^ can induce a prolonged torpor-like state, with QRFP neurons (Q neurons, for simplicity) specifically contributing to an intensive hypothermic condition known as Q neuron-induced hypometabolic/hypothermic state (QIH)^16^. In addition, cold-sensitive POA neurons expressing bombesin-like receptor 3 (BSR3) are involved in inducing thermogenesis^18^. The identification of these molecularly defined neural populations facilitates the monitoring of their activities during thermal challenges^13, 18^ and torpor periods^17^, manipulation of their functions^11, 15–19^, and mapping of their input/output neural circuitry^11, 16^. Regarding the thermosensory input to the POA, extensive research has revealed that ascending cold and warm signals are mediated by two anatomically distinct subdivisions of the lateral parabrachial nuclei (LPB) in the brainstem: cold signals in the external lateral LPB (LPBel) and warm signals in the dorsal LPB (LPBd)^20–24^. Although the detailed circuit mechanisms remain incompletely understood, these signals are thought to be integrated into POA neurons.

POA neurons project to downstream neural circuits extending from the dorsomedial hypothalamus (DMH) to the medullary RPa^2^. DMH neurons are activated by cold stimuli *in vivo*^25^, and the genetic activation of DMH neurons expressing vesicular glutamate transporter 2 (vGluT2), LeptR, and BRS3 induces thermogenesis^13, 26, 27^. RPa neurons receive input from the DMH, and optogenetic activation of this pathway induces BAT thermogenesis^26, 28, 29^. Exposure to cold or pyrogens induces c-Fos expression, used as a proxy for neural activation, in RPa neurons expressing *vGluT3*^30, 31^. These *vGluT3*-expressing (+) neurons project axons to the spinal cord to modulate sympathetic outflow to the BAT. Pharmacological inhibition of the RPa impairs both BAT and shivering thermogenesis^28, 30, 32^.

In contrast to the substantial progress made in the characterization of thermoregulatory neurons in the POA and DMH, research on medullary thermoregulatory systems has lagged behind. Although recent seminal studies have utilized *vGluT3-Cre* mice to genetically access *vGluT3+* neurons in the RPa, demonstrating their capacity to drive lipolysis^33^ and thermogenesis^34^ through chemogenetic activation, these studies have not elucidated the precise neural activity dynamics of these neurons or mapped their input/output neural circuit organization. To address this gap and facilitate Cre-based neural manipulations, we generated *vGluT3-Flpo* mice, which allow fiber photometry monitoring^35^ of *vGluT3*+ neuron activity in conjunction with Cre-mediated targeted modulation of upstream Q neurons^16^. Additionally, we utilized rabies virus-mediated monosynaptic tracing^36^ along with axonal projection mapping to comprehensively characterize the inputs and outputs of *vGluT3+* neurons in the RPa, shedding light on their activity dynamics, functions, and neural circuitry in the context of thermoregulation in mice.

## Results

### Generation of *vGluT3-Flpo* knock-in mice

To genetically target *vGluT3*+ neurons, *vGluT3-Flpo* knock-in mice were generated using CRISPR-mediated genome editing (Fig. S1)^37^. To test the efficiency and specificity of Flpo activity, we injected an Flpo-dependent adeno-associated virus (AAV) vector encoding GCaMP6s^38^ into the RPa of *vGluT3-Flpo* mice (Fig. 1A). Brain sections were stained with a *vGluT3* RNA probe by *in situ* hybridization (ISH), combined with immunostaining, for green fluorescent protein (GFP) (Fig. 1B). We found that 95.4 ± 2.4% (mean ± standard deviation) of GCaMP6s+ neurons expressed *vGluT3* and the efficiency of viral targeting was approximately 40% (Fig. 1C). Sections were also stained with tryptophan hydroxylase 2 (TPH2), a marker for serotonergic neurons within the RPa, which are implicated in regulating the respiratory systems^39–41^ and exhibit an inverse correlation with *vGluT3*^42^ mRNA expression levels. We found that only 2.3 ± 2.8% of GCaMP6s+ neurons expressed TPH2. These data indicate that Flpo expression is specific to a subpopulation of non-serotonergic *vGluT3*+ neurons within the RPa (hereafter referred to as RPa-*vGluT3* neurons).

**Figure 1:**
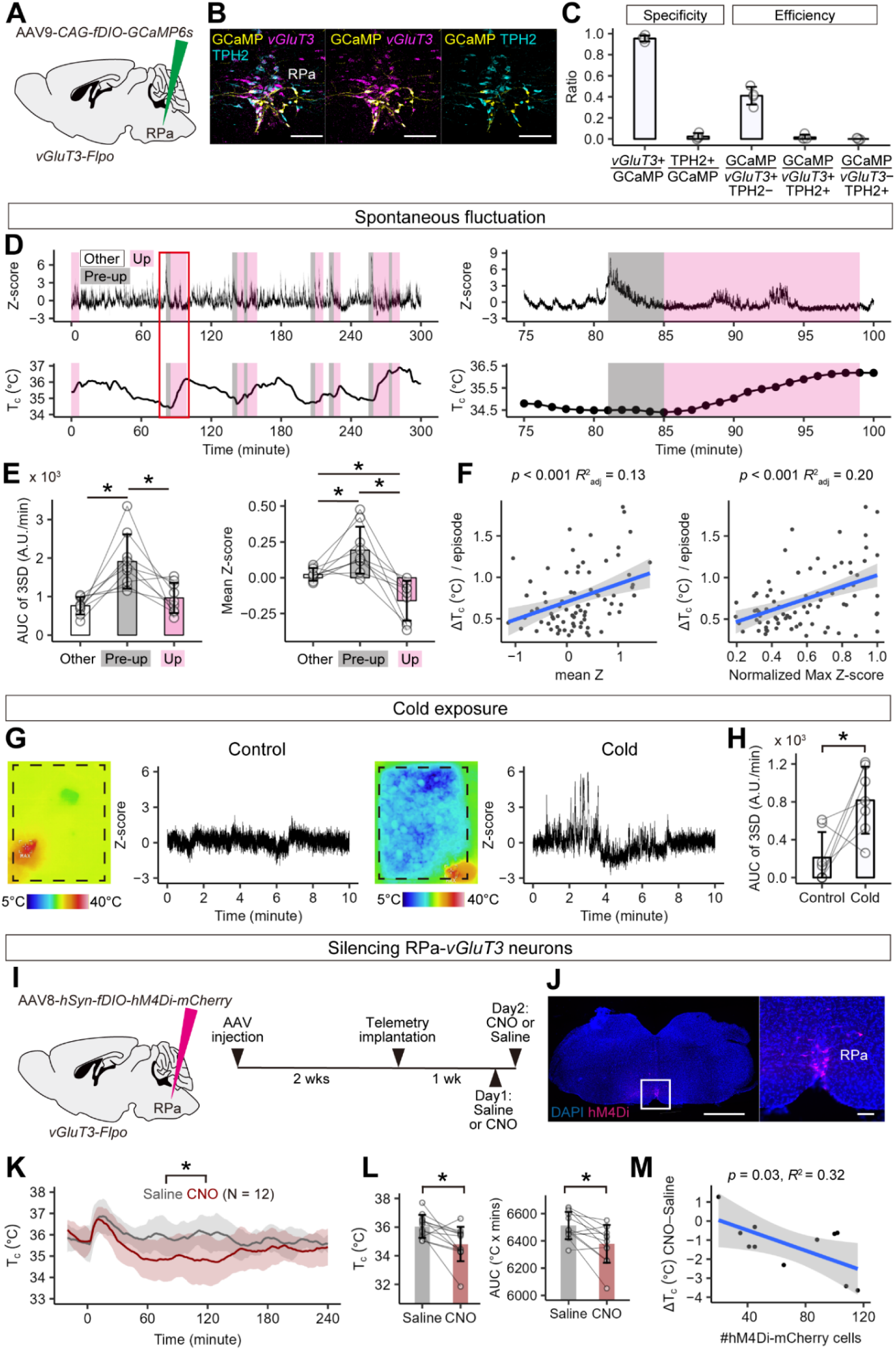
Fiber photometry-based Ca^2+^ imaging of RPa-*vGluT3* neurons. (A) Schematic of the experiment. (B) Representative coronal section of the RPa showing *vGluT3* (magenta), GCaMP6s (yellow), and TPH2 (cyan) expression. Scale bars, 100 µm. (C) Targeting specificity (*vGluT3*+/GCaMP6s+ and TPH2+/GCaMP6s+) and targeting efficiency (GCaMP6s+/*vGluT3*+TPH2−, GCaMP6s+/*vGluT3*+TPH2+, and GCaMP6s+/*vGluT3*−TPH2+). N = 4 mice. (D) Representative photometry (top) and T_c_ (bottom) traces. Gray and magenta shading represent the pre-up and up phases, respectively. Right panels show 25-min magnified views of the red-boxed region. (E) AUC of 3SD signals (left) and mean Z-score (right). Friedman rank sum test with pairwise comparisons using the Wilcoxon signed-rank sum test with Bonferroni correction, \**p* < 0.05, N = 9 mice. (F) Correlation between the mean Z-score or normalized maximum Z-score during the pre-up phase (left) and T_c_ increase per episode. n = 86 episodes from 9 mice. Adjusted coefficient of determination (R^2^) is shown; p-value calculated using a t-test under the null hypothesis of no correlation. (G) Representative photometry traces under control room temperature (left) and cold exposure (right) conditions. (H) AUC of 3SD signals. \**p* < 0.05 by exact Wilcoxon signed-rank sum test. N = 7 mice. (I) Schematic of the experimental procedure (left) and timeline (right) for chemogenetic inhibition of RPa-*vGluT3* neurons. (J) Representative coronal sections showing hM4Di (magenta) expression in the RPa, with DAPI nuclear counterstaining (blue). Scale bars, 1 mm (left) and 100 µm (right). (K) Group mean traces of T_c_ in hM4Di+ mice following administration of saline (gray) or CNO (red) at time 0. Repeated-measures two-way ANOVA shows significant solution (*p* < 0.01), time course (*p* < 0.01), and interaction (*p* < 0.01) effects. (L) T_c_ at 120 min following saline or CNO injection (left) and AUC for T_c_ from 0 to 240 min (right). *, *p* < 0.05 by Wilcoxon signed-rank sum test. N = 12. (M) Correlation between the number of hM4Di-mCherry+ cells and the change in T_c_ at 120 min following saline or CNO injection. Adjusted coefficient of determination (R^2^) is shown, with the *p*-value calculated using a *t*-test under the null hypothesis of no correlation. Error bars indicate standard deviation (SD). For additional data, see Figs. S1–S3.

### Phasic activity of RPa-*vGluT3* neurons during thermogenesis

To visualize the activity of RPa-*vGluT3* neurons, we used fiber photometry-based *in vivo* Ca^2+^ imaging^35^. We injected AAV*-fDIO-GCaMP6s* into the RPa and placed optical fibers over the RPa of *vGluT3-Flpo* mice (Fig. S2A). Post hoc histology confirmed the expression of GCaMP6s and the locations of the optic fibers (Fig. S2B). We used a telemetry system to monitor T_c_ in freely moving mice (Fig. 1D). The representative trace in Fig. 1D shows the phasic elevations in photometric signals. To correlate these signals with T_c_ variations, we defined the “up phase” or “down phase” as periods during which the T_c_ continuously increased or decreased for ≥5 min, with a change of at least 0.3°C. When analyzing the area under the curve (AUC) of the photometric signals showing peak amplitudes exceeding +3 standard deviations (hereafter referred to as 3SD signals for simplicity), no differences were found among the up, down, and other phases (Fig. S2C and S2D). Conversely, when the pre-up phase was defined as the 4-min window preceding the onset of the up phase, both the AUC of the 3SD signals and the mean Z-scored signal intensity were significantly larger during the pre-up phase than during the up and other phases (Fig. 1E). Furthermore, a significant positive correlation was found between the mean or maximum Z-score signal intensity during the pre-up phase and the increase in T_c_ (Fig. 1F). These findings suggest that RPa-*vGluT3* neuron activity is phasically upregulated before a spontaneous rise in T_c_ and that this activity may influence the magnitude of the T_c_ increase.

Next, we investigated whether cold exposure induces Ca^2+^ responses in RPa-*vGluT3* neurons. To address this, the floor temperature was lowered to below 15°C by placing an ice pack beneath the home cage, while a room-temperature pack served as the control (Fig. 1G). Cooling the floor significantly increased the AUC of the 3SD signals (Fig. 1H), supporting the notion that cold sensory signals activate RPa-*vGluT3* neurons to drive thermogenesis.

### Silencing RPa-*vGluT3* neurons decreases T_c_ and delays recovery from QIH

To assess whether the basal activity of RPa-*vGluT3* neurons is required to maintain T_c_, we utilized a chemogenetic inhibition strategy. AAV8-*hSyn-fDIO-hM4Di-mCherry* was bilaterally injected into the RPa, followed by the implantation of a telemetry probe (Fig. 1I). Post hoc histological analysis confirmed the localized expression of hM4Di within the RPa (Fig. 1J). Intraperitoneal injections of either saline or clozapine-N-oxide (CNO) were administered to the mice on consecutive days. Administration of CNO significantly reduced both T_c_ and BAT surface temperature (T_BAT_) (Fig. 1K, 1 L, Fig. S3A, and S3B). Notably, the extent of CNO-induced hypothermia was significantly correlated with the number of hM4Di-mCherry+ cells in the RPa, supporting the role of these neurons in maintaining T_c_ (Fig. 1M and Fig. S3C).

We further examined whether RPa-*vGluT3* neurons are required for cold tolerance. First, we assessed the effects of their inhibition during acute cold exposure. Intraperitoneal injections of either saline or CNO were administered to the mice, and 30 min later, their home cages were placed on an ice pack. CNO administration to inhibit RPa-*vGluT3* neurons slightly but significantly reduced T_c_ (Fig. S3D), further supporting the idea that RPa-*vGluT3* neurons are needed to drive cold-defense thermogenesis. T_BAT_ remained unchanged under this condition (Fig. S3E), implying that cold-induced thermogenesis in BAT can be activated through sympathetic-independent mechanisms^43^.

Recent studies have demonstrated that the activation of specific neuronal populations within the POA induces hypothermic states, likely through a reduction in the T_c_ set point^10, 12, 13, 15–17^. For instance, activation of Q neurons in the anteroventral periventricular nucleus (AVPe), a POA subregion, has been shown to induce QIH^16^. To investigate the activity dynamics of RPa-*vGluT3* neurons during QIH, we generated *vGluT3-Flpo*; *Qrfp-iCre*^16^ double-heterozygous mice. AAV5*-EF1a-DIO-hChR2-eYFP* was injected into the AVPe, whereas AAV9-*CAG-fDIO-GCaMP6s* was delivered to the RPa, followed by the implantation of optical fibers above these regions (Fig. 2A). Post hoc histological analysis confirmed the expression of ChR2-YFP and GCaMP6s, as well as proper fiber placement (Fig. 2B). One week after telemetry implantation, we monitored Ca^2+^ responses in RPa-*vGluT3* neurons using fiber photometry while inducing hypothermia via laser stimulation of Q neurons for 3 h (Fig. 2C). Before QIH induction, the photometric signals exhibited high variability. During QIH, RPa-*vGluT3* neuronal activity was markedly suppressed. Notably, upon termination of hChR2 stimulation, the photometric signals resumed. The AUC of the 3SD signals was significantly reduced during QIH, with no differences observed before and after QIH (Fig. 2D). A detailed temporal analysis revealed that the decline in T_c_ preceded a reduction in RPa-*vGluT3* neuronal activity (Fig. 2E and 2F), whereas the reactivation of RPa-*vGluT3* neurons occurred prior to T_c_ recovery (Fig. 2G and 2H). These data indicate that RPa-*vGluT3* neuron activity is dynamically regulated throughout the QIH process.

**Figure 2:**
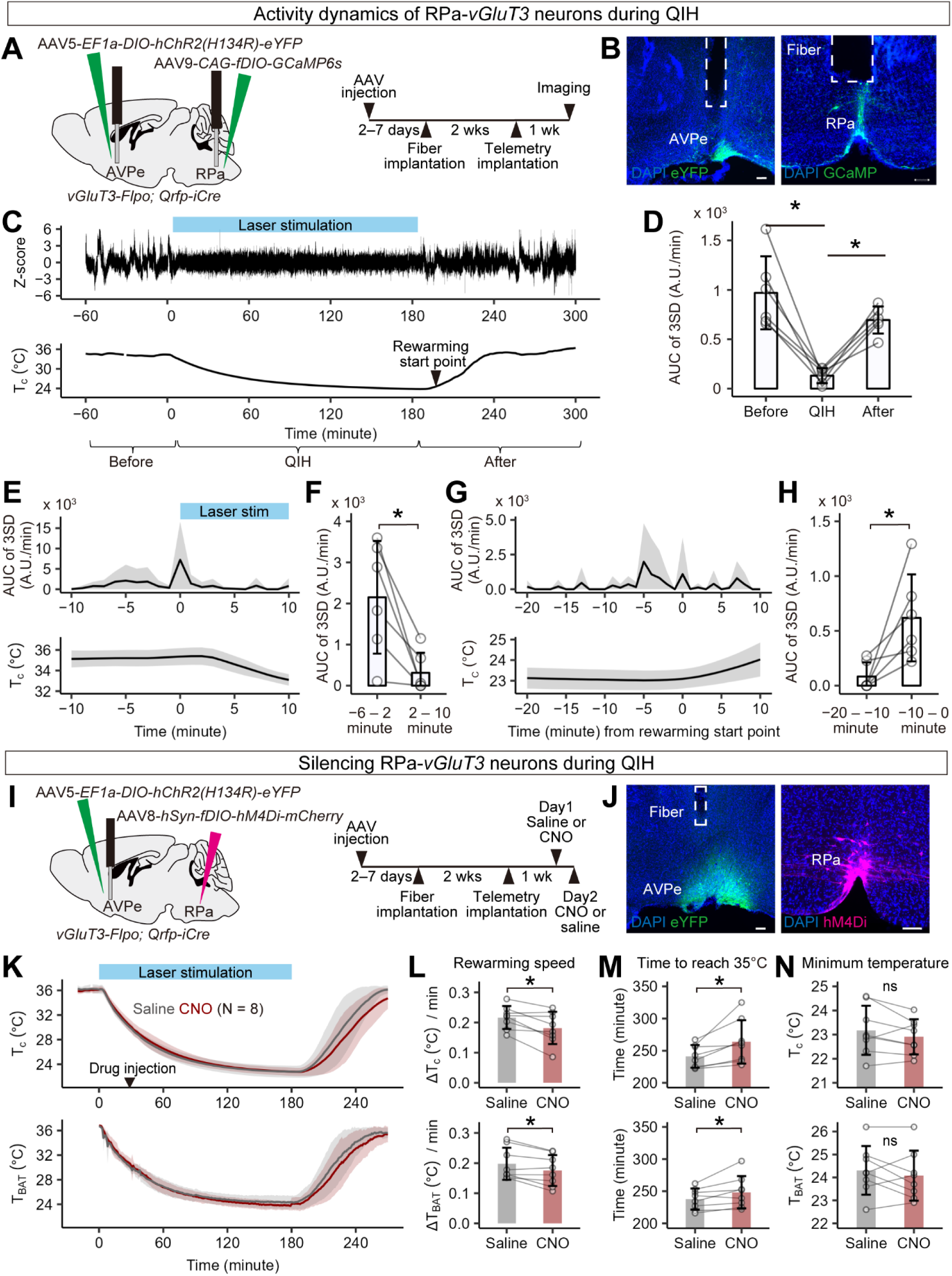
Silencing RPa-*vGluT3* neurons delays recovery from QIH. (A) Schematic of the experimental procedure (left) and timeline (right) for fiber photometry of RPa-*vGluT3* neurons combined with optogenetically induced QIH. (B) Representative coronal sections showing ChR2-eYFP (green) expression in the AVPe (left) and GCaMP (green) expression in the RPa (right), with DAPI nuclear counterstaining (blue). Scale bar, 100 µm. (C) Representative photometry (top) and T_c_ (bottom) traces during QIH. Blue bar indicates the period of laser stimulation. Arrow indicates T_c_ rise point. (D) AUC of 3SD signals. *, *p* < 0.05 by pairwise t-tests with Bonferroni correction. N = 6 mice. (E, G) 3SD signal durations (top) and change in T_c_ (bottom) relative to QIH induction (E) or the T_c_ rise point following cessation of laser stimulation (G). Bold lines indicate group means; gray shading represents SD. (F, H) AUC of 3SD signals within designated 4- or 10-min time windows. *, *p* < 0.05 by Wilcoxon signed-rank sum test. N = 6 mice. (I) Schematic of virus injection (left) and experimental timeline for chemogenetic inhibition of RPa-*vGluT3* neurons during QIH (right). (J) Representative coronal sections showing ChR2-eYFP (green) expression in the AVPe (left) and hM4Di-mCherry (magenta) expression in the RPa (right), with DAPI staining (blue). Scale bar, 100 µm. (K) T_c_ (top) and T_BAT_ (bottom) traces following laser stimulation (blue bar). CNO (red) or saline (gray) was administered 2.5 h before laser cessation. Lines represent group means; shading denotes SD. N = 8 each. (L–N) Rewarming speed (L), latency to reach 35°C (M), and minimum temperature (N) for T_c_ (top) or T_BAT_ (bottom) after saline or CNO injection. * *p* < 0.05, Wilcoxon signed-rank sum test. N = 8 each. Error bars indicate SD. For more data, see Fig. S3.

To further investigate the role of RPa-*vGluT3* neurons in rewarming after QIH, we chemogenetically inhibited these neurons using hM4Di in combination with optogenetically induced QIH (Fig. 2I and 2J). If RPa-*vGluT3* neurons facilitate thermogenesis during rewarming, then their inactivation delays recovery. Following AAV injection and fiber/telemetry implantation, intraperitoneal injections of either saline or CNO were administered to the mice 30 min after the onset of laser stimulation. Laser illumination was terminated 2.5 h later, and the subsequent recovery of T_c_ was assessed. CNO administration significantly slowed the rewarming rate of both T_c_ and T_BAT_ (Fig. 2K and 2L), and prolonged the latency for T_c_ and T_BAT_ to reach 35°C (Fig. 2M). Conversely, the minimum T_c_ attained during QIH remained unchanged in the saline- and CNO-injected groups (Fig. 2N). Notably, the CNO-induced delay in T_c_ recovery was positively correlated with the number of hM4Di+ neurons in the RPa (Figs. S3F and S3G); CNO administration did not affect rewarming in mice with limited hM4Di expression (Fig. S3H–S3K). These data demonstrate that RPa-*vGluT3* neurons facilitate thermogenesis during recovery from QIH.

### Input and output architecture of RPa-*vGluT3* neurons

Previous studies have suggested that RPa-*vGluT3* neurons receive excitatory inputs from glutamatergic neurons in the DMH^26^ and project to sympathetic preganglionic neurons within the thoracic spinal cord^31, 44^. However, the precise organization of their afferent and efferent connectivity remains unclear. To systematically map the input connections of RPa-*vGluT3* neurons, we utilized rabies virus (RV)- mediated retrograde trans-synaptic tracing^36^ (Fig. 3A). Starter neurons, defined by the overlap of TVA-mCherry and RV-derived nuclear GFP (nGFP), were primarily located in the RPa (Fig. 3B). Retrogradely labeled input neurons were distributed across various regions, including the DMH and ventrolateral periaqueductal gray (vlPAG) (Fig. 3C–3F). A negative control experiment, in which AAV*-fDIO-RG* was omitted, showed nGFP+ mCherry− cells as nonspecific RV labeling^36^, with the vast majority located near the injection site, particularly within the gigantocellular reticular nucleus (Gi) (Fig. S4). To ensure the accuracy of our analysis, regions with substantial nonspecific labeling were excluded. We further characterized the cell types of the nGFP-labeled input neurons in the DMH and vlPAG by immunostaining for RV-nGFP combined with ISH using excitatory (*vGluT2*) or inhibitory (*vGat*) neuronal markers. We found that 83.6 ± 10.5% of the input neurons in the DMH were *vGluT2+*, while 92.1 ± 0.6% of input neurons in the vlPAG were *vGat+* (Fig. 3G–3I). While glutamatergic neurons in the DMH in promoting thermogenesis is well established^26, 28, 29^, inhibitory neurons in the vlPAG and the adjacent dorsal raphe nucleus have been reported to suppress thermogenesis^45^. These data indicate that RPa-*vGluT3* neurons receive prominent excitatory inputs from the DMH while integrating diverse brainstem inputs, particularly inhibitory inputs from the vlPAG.

**Figure 3:**
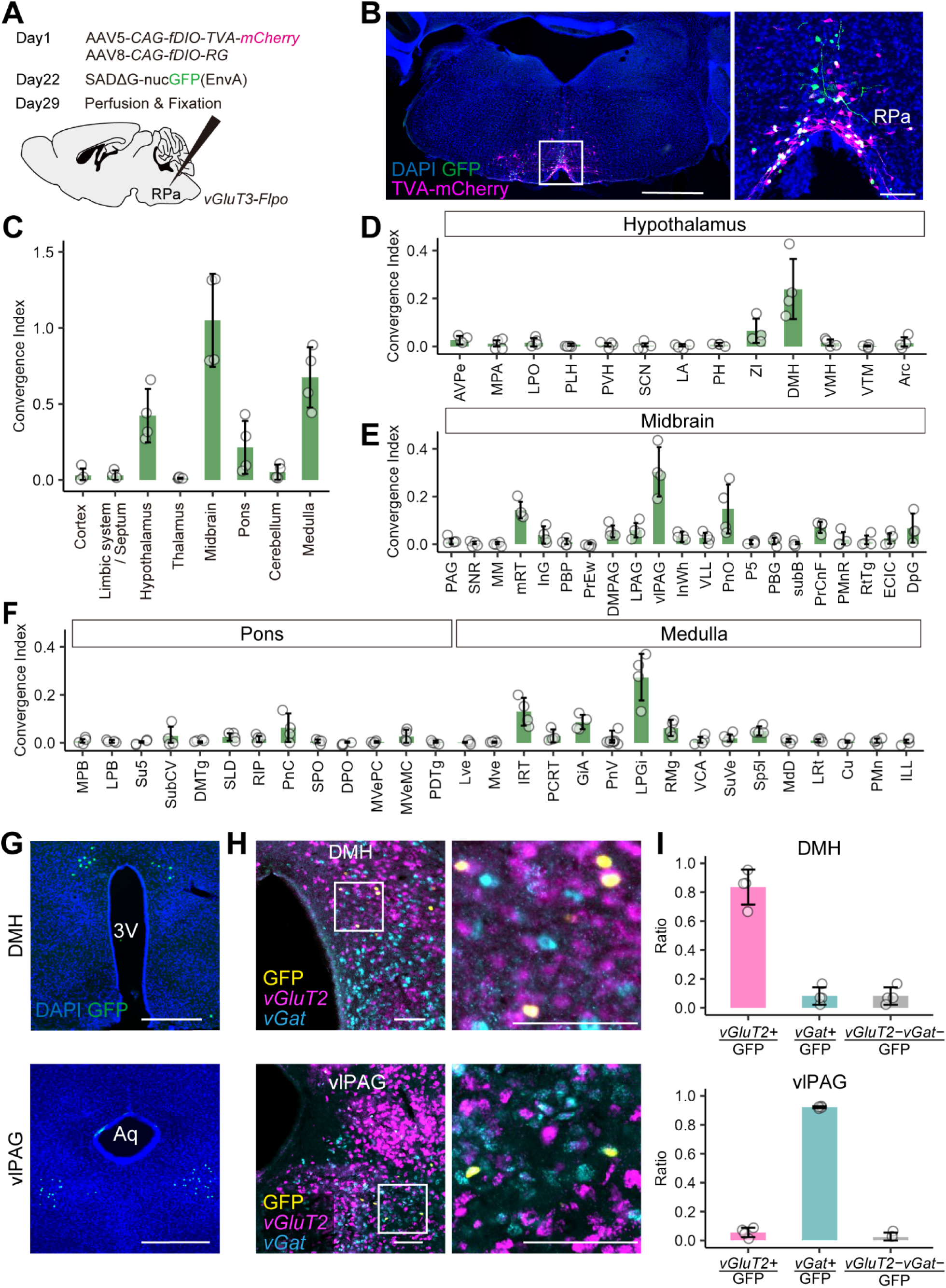
Input map to RPa-*vGluT3* neurons. (A) Experimental procedure for RV-mediated trans-synaptic tracing. (B) Representative coronal section of the RPa showing GFP expression from RV (green) and TVA-mCherry labeling from AAV (magenta) with DAPI nuclear counterstaining (blue). Right panels show a magnified view of the white-boxed region. Scale bars, 1 mm (left) and 100 µm (right). (C) Convergence index (number of input cells normalized to the number of starter cells) across eight broader anatomical categories. N = 4 mice. (D–F) Convergence indices for various brain regions in the hypothalamus (D), midbrain (E), and pons and medulla (F). N = 4 mice. (G) Representative coronal sections showing labeled input neurons in the DMH (left) and vlPAG (right). Scale bars, 500 µm. (H) Representative coronal sections of the DMH (left) and vlPAG (right) showing *vGluT2* (magenta) and *vGat* (cyan) mRNA expression along with anti-GFP immunostaining (yellow). Bottom panels show magnified views of the corresponding white-boxed regions. Scale bar, 100 µm. (I) Fraction of *vGluT2+*, *vGat+*, and dual-negative neurons among GFP-labeled input neurons in the DMH (left) and vlPAG (right). N = 4 mice. Error bars indicate SD. For more details, see Fig. S4. For brain region abbreviations, see Table S1.

To delineate the efferent projections of RPa-*vGluT3* neurons, we injected AAV-*EF1a*-*fDIO-mCherry* into the RPa of *vGluT3-Flpo* mice and mapped the distribution of mCherry-positive axons in both the brain and spinal cord (Fig. 4A and 4B). Dense mCherry+ axonal projections were observed in the intermediolateral nucleus (IML) of the upper thoracic spinal cord (Fig. 4C and 4E), consistent with previous studies^31, 44^. mCherry+ axons were detected in various brainstem regions (Fig. 4C and 4D). Notably, projections were observed in the ventral brainstem nuclei, such as the supratrigeminal nucleus (Su5), intermediate reticular formation (IRT), and ventral (MdV) and dorsal (MdD) medullary reticular nuclei, which are regions associated with somatic premotor control and the regulation of intrinsic behaviors^46–49^. In the dorsal brainstem, we observed the axonal projections of RPa-*vGluT3* neurons in the LPB, a region implicated in the transmission of cold sensory information to the hypothalamus and limbic structures^22, 23^. These findings demonstrate previously uncharacterized efferent pathways of RPa-*vGluT3* neurons, suggesting a broader functional role beyond their proposed function as sympathetic premotor neurons mediating non-shivering thermogenesis^31^.

**Figure 4:**
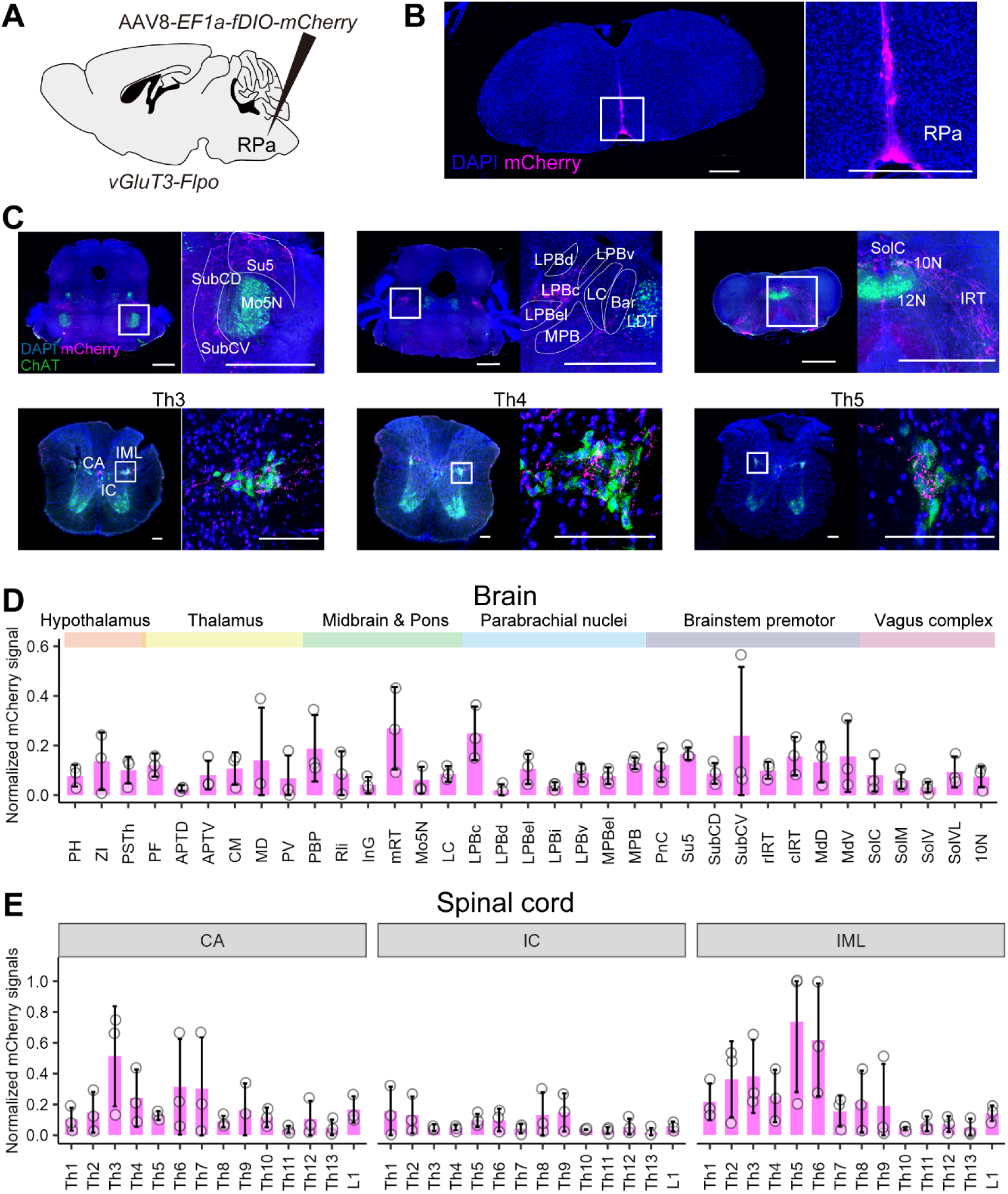
Axonal projections of RPa-*vGluT3* neurons. (A) Schematic of AAV injection. (B) Representative coronal section of the RPa showing mCherry expression. Right panel shows a magnified view of the white-boxed area. Scale bar, 500 µm. (C) Representative coronal sections of the brainstem (top) and spinal cord areas (bottom). Right images show magnified views of the white-boxed regions. Green, anti-ChAT immunostaining; magenta, anti-mCherry immunostaining; blue, DAPI nuclear staining. Scale bars, 1 mm (top) and 100 µm (bottom). Th, thoracic segment. (D, E) Quantification of mCherry+ axons in the brain (D) and spinal cord (E). Axonal projections were quantified and normalized to the maximum value per mouse. N = 3 mice. Error bars indicate SD. For brain region abbreviations, see Table S1.

### RPa-*vGluT3* neurons regulate multiple thermo-effector pathways

To examine whether RPa-*vGluT3* neurons regulate multiple thermogenic responses, we targeted hChR2 in these neurons using a multimodal physiological monitoring approach. T_c_ was measured via telemetry, T_BAT_ was assessed using infrared thermography, and thermogenic shivering was evaluated using electromyography (EMG) recordings of nuchal muscle activity (Fig. 5A and 5B). Mice were connected to a laser system and subjected to optogenetic stimulation for 10 min. This procedure significantly increased both T_c_ and T_BAT_ in hChR2+ mice compared to mCherry+ controls (Fig. 5C and 5D), confirming that RPa-*vGluT3* neurons promote thermogenesis via BAT^34^. We also confirmed that the chemogenetic activation of RPa-*vGluT3* neurons led to an increase in T_c_ (Fig. S5). In addition to thermogenesis, the hChR2+ mice exhibited cervicothoracic piloerection during photostimulation (Fig. 5E and Movie S1), suggesting the recruitment of additional sympathetic effectors. Consistent with this view, optogenetic activation of RPa-*vGluT3* neurons significantly induced c-Fos expression, a proxy for neural activation, in postganglionic neurons projecting to both the BAT and cervical skin (Fig. S6A–S6C). Thus, RPa-*vGluT3* neurons engage in multiple sympathetic outputs to drive non-shivering thermogenesis via BAT and heat retention via piloerection.

**Figure 5:**
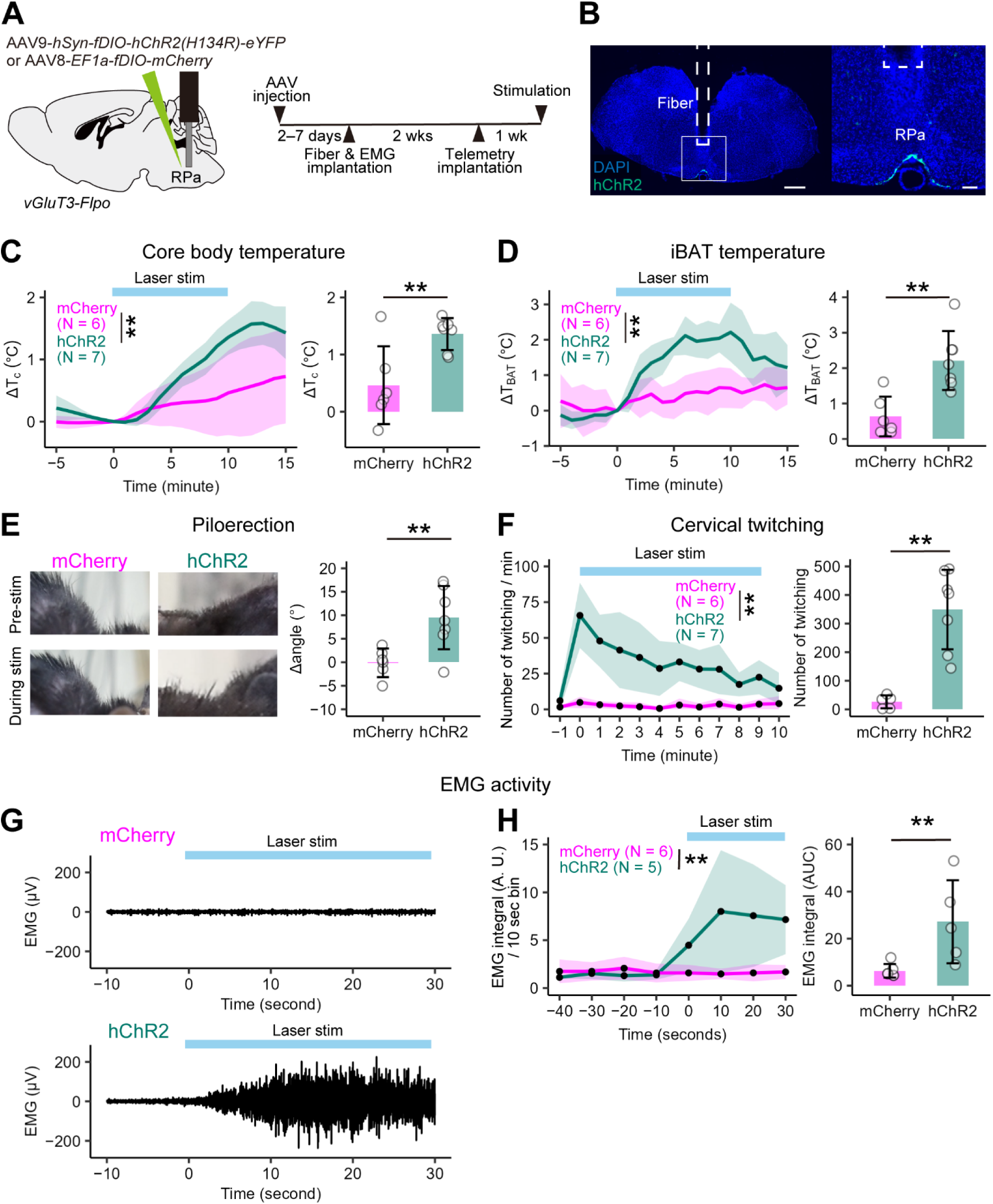
Optogenetic activation of RPa-*vGluT3* neurons induces multiple thermogenic responses. (A) Schematic of the experimental procedures (left) and timeline (right). (B) Representative coronal sections of the RPa showing ChR2-eYFP (green) expression with DAPI nuclear counterstaining (blue). Right panel shows the magnified view of the white-boxed region. Scale bars, 500 µm (left) and 100 µm (right). (C, D) Change in T_c_ (C) and T_BAT_ (D) following laser stimulation (blue shading). Green traces denote data from hChR2+ mice; magenta traces represent mCherry+ control mice. Lines represent group means; shading denotes SD. Two-way ANOVA: AAV effect, *p* < 0.01; time effect, *p* < 0.01; interaction effect, *p* < 0.01. Right panels show ΔT_c_ (C) and ΔT_BAT_ (D) at 10 min post-stimulation. **, *p* < 0.01 by Wilcoxon rank sum test. N = 6 for the mCherry group; N = 7 for hChR2 group. (E) Representative images of cervicothoracic piloerection before and during laser stimulation in mCherry+ control (left) and hChR2+ (right) mice. Right panel shows quantification of hair angle changes. *, *p* < 0.05 by two-sided unpaired *t*-test. N = 6 for the mCherry group and N = 7 for hChR2 group. (F) Quantification of cervical muscle twitching. Two-way ANOVA: AAV effect, *p* < 0.01; time effect, *p* < 0.01; interaction effect, *p* < 0.01. Right panel shows total number of twitch episodes during the laser stimulation. *, *p* < 0.05 by Wilcoxon rank sum test. N = 6 for the mCherry group and N = 7 for the hChR2 group. (G) Representative EMG traces from the mCherry (upper) and hChR2 (lower) groups during laser stimulation (blue shading) under isoflurane anesthesia. (H) Quantification of EMG integrals per 10 s bin during laser stimulation. Magenta (mCherry) and green (hChR2) lines represent group means; shading denotes SD. Two-way ANOVA: AAV effect, *p* < 0.01; time effect, *p* < 0.01; interaction effect, *p* < 0.01. Right panel shows EMG integrals during laser stimulation. *, *p* < 0.05 by two-sided unpaired *t*-test. N = 6 for the mCherry group and N = 5 for the hChR2 group. For further data, see Figs. S5 and S6.

In addition to sympathetic output, RPa-*vGluT3* neuron stimulation triggered shivering-like twitches in the cervical muscles (Fig. 5F and Movie S1). EMG recordings revealed a significant increase in integrated neck muscle activity following optogenetic stimulation of RPa-*vGluT3* neurons (Fig. 5G, 5H), implicating activation of the somatic thermo-effector pathway independent of the sympathetic nervous system. In addition, optical stimulation increased locomotor activity in hChR2+ mice (Fig. S6D).

We hypothesized that RPa-*vGluT3* neurons mediate thermogenic shivering via descending projections to brainstem premotor areas (Fig. 4D), particularly the caudal IRT (cIRT), a known regulator of neck muscle movement^47, 50^. To test this hypothesis, we injected AAV9-*EF1a-fDIO- hChR2-eYFP* into the RPa and AAV8-*CaMKIIa-hM4D(Gi)-mCherry* into the cIRT, followed by optical fiber implantation above the RPa (Fig. 6A, 6B). Optogenetic stimulation consistently elevated T_c_ and T_BAT_ and triggered piloerection, regardless of CNO administration (Fig. 6C–6E). In contrast, CNO-mediated inhibition of cIRT neurons tended to reduce cervical muscle twitches (Fig. 6F) and significantly suppressed EMG activity (Fig. 6G, 6H). Thus, the shivering-like activity of nuchal muscles triggered by RPa-*vGluT3* neurons is at least partly mediated by cIRT neurons.

**Figure 6:**
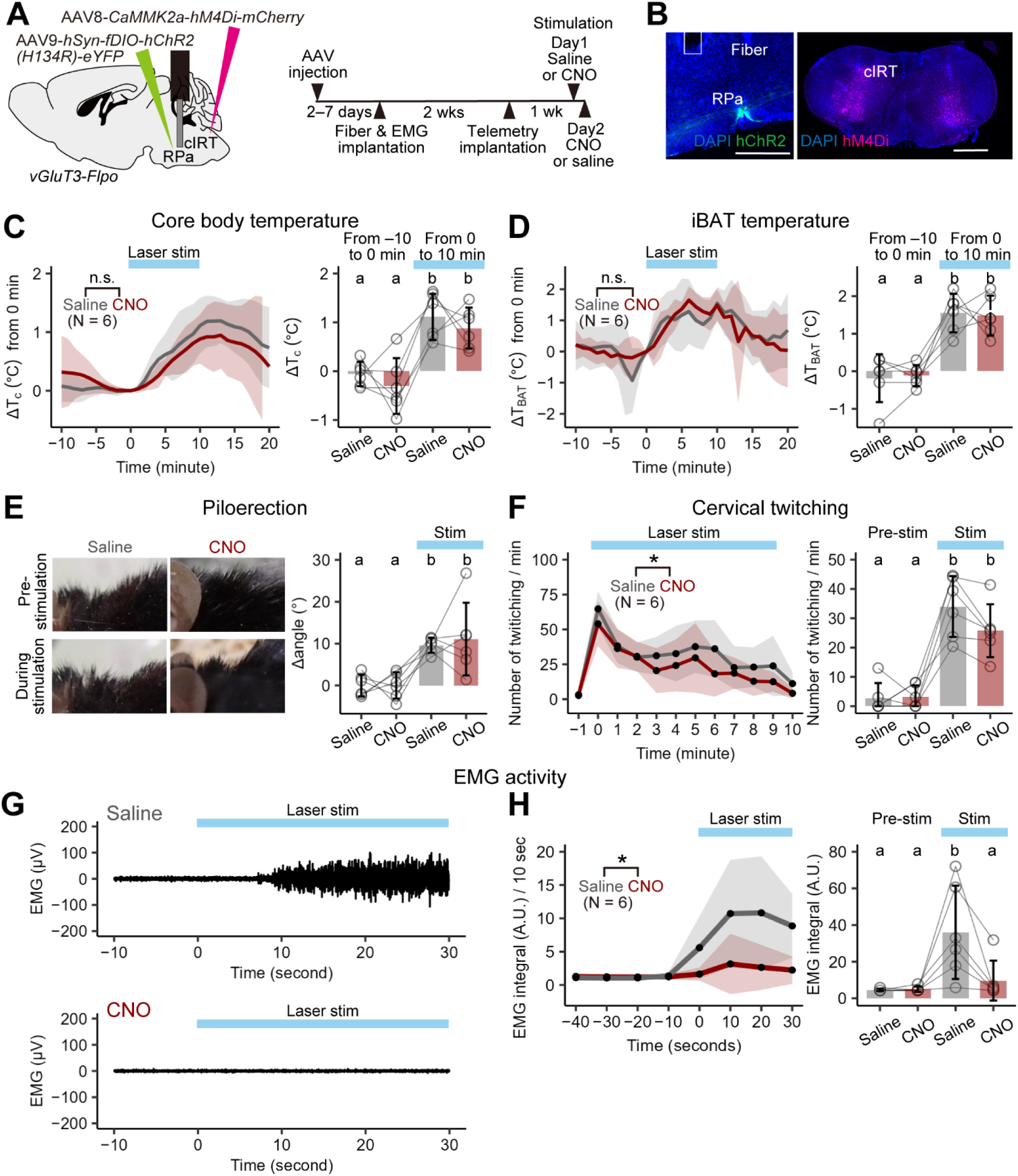
Optogenetic activation of RPa-*vGluT3* neurons induces shivering via brainstem premotor neurons. (A) Schematic of the experimental procedures (left) and timeline (right). (B) Representative coronal sections. Upper panel shows the RPa with ChR2-eYFP (green) expression and DAPI nuclear counterstaining (blue). The bottom panel shows hM4Di-mCherry (magenta) expression in the cIRT with DAPI (blue). Scale bars, 500 µm. (C, D) Change in T_c_ (C) and T_BAT_ (D) following laser stimulation (blue shading). CNO (red) or saline (gray) was administered 45–60 min before laser stimulation. Lines represent group means; shading denotes SD. N = 6 each. Repeated-measures two-way ANOVA: Drug effect, n.s.; time effect, *p* < 0.01; interaction effect, n.s. Right panels show ΔT_c_ (C) and ΔT_BAT_ (D) during the 10-min before (−10 to 0 min) and during (0 to 10 min) laser stimulation. (E) Representative images of cervicothoracic piloerection before and during laser stimulation in saline- (left) and CNO- (right) treated mouse. Right panel shows quantification of hair angle changes. (F) Quantification of cervical muscle twitching. Left panel shows the total number of twitch episodes during laser stimulation. Repeated-measures two-way ANOVA: Drug effect, *p* < 0.05; time effect, *p* < 0.01; interaction effect, n.s. Right panel shows total number of twitch episodes per minute before and during the laser stimulation. (G) Representative EMG traces from saline- (upper) and CNO- (lower) treated mice during laser stimulation (blue shading) under anesthesia. (H) Quantification of EMG integrals per 10 s bin during laser stimulation. Black (saline) and magenta (CNO) lines represent group means; shading denotes SD. Repeated-measures two-way ANOVA: Drug effect, *p* < 0.05.; time effect, *p* < 0.01; interaction effect, *p* < 0.01. Right panel shows EMG integrals for 40 s before and during laser stimulation. In all panels, different letters (a, b) indicate significant differences at *p* < 0.05, as determined by one-way repeated-measures ANOVA followed by Tukey–Kramer post hoc test. N = 6 per group.

Collectively, these data support the idea that RPa-*vGluT3* neurons recruit multiple thermo-effector pathways, including sympathetic outputs to BAT and skin, and descending projections to somatic premotor regions, such as the cIRT, to drive coordinated thermogenic responses.

## Discussion

In the present study, we generated *vGluT3-Flpo* mice, facilitating targeted viral-genetic approaches to monitor the activity dynamics and map the input-output architecture of RPa-*vGluT3* neurons (Fig. S7). Below, we discuss the biological insights derived from this study and its limitations.

### Activity dynamics of RPa-*vGluT3* neurons

While classical c-Fos mapping has implicated RPa-*vGluT3* neurons in cold exposure and PGE_2_-induced thermoregulatory responses^31^, the technique’s limitation of temporal resolution has precluded precise characterization of their activity dynamics. Our fiber photometry data demonstrated that RPa-*vGluT3* neurons exhibited sharp phasic activity immediately preceding a spontaneous T_c_ increase (Fig. 1D), during cold exposure (Fig. 1G), and before recovery from QIH (Fig. 2C). Notably, there was a consistent delay of several minutes between peak neural activity and the onset of T_c_ elevation. This temporal gap likely reflects the time required to recruit thermogenic and heat-retention systems (e.g., BAT activation, shivering, and piloerection) and the intrinsic thermal inertia of the body. By the time T_c_ begins to rise (the up phase in Fig. 1), RPa-*vGluT3* neuron activity has already declined, suggesting the presence of a rapid shutoff mechanism that may prevent overshooting of T_c_. Taken together, our data highlight the phasic nature of RPa-*vGluT3* neuron activity. Given that chemogenetic silencing of these neurons reduced both the baseline T_c_ and cold-evoked thermogenesis (Fig. 1I–1M and Fig. S3A–S3E), their phasic activity seems to be critical for maintaining thermal homeostasis.

During QIH, the metabolic demand, as indicated by oxygen consumption, is markedly reduced, leading to a subsequent drop in T_c_ ^16^. This presumed decrease in the central T_c_ set point is thought to suppress medullary thermogenic systems, preventing BAT activity and shivering even when T_c_ falls to 25°C. However, this hypothesis has not yet been tested experimentally. Our data provide direct evidence of a pronounced suppression of RPa-*vGluT3* neuron activity, aligned with the optogenetic activation of Q neurons in the AVPe (Fig. 2C). Furthermore, the chemogenetic silencing of RPa-*vGluT3* neurons resulted in a significant delay in T_c_ recovery following QIH, reinforcing their role in thermogenic reactivation. The neural circuit mechanisms by which Q neurons exert this rapid and potent inhibitory effect on RPa-*vGluT3* neurons remain unclear and require further investigation.

### Neural circuit organizations of RPa-*vGluT3* neurons

The prevailing model of the medullary thermogenic system assumes that it acts as a relay, transmitting thermal commands from the preoptic and hypothalamic regions to downstream sympathetic circuits that drive BAT activation^1, 2^. Our data support this framework by providing anatomical evidence that RPa-*vGluT3* neurons receive direct excitatory synaptic inputs from DMH neurons (Fig. 3H), a key hypothalamic output, and project to the IML of the thoracic spinal cord for sympathetic activation of BAT thermogenesis (Fig. 4E and Fig. S6). However, our data also demonstrate that RPa-*vGluT3* neurons play a broader role in thermoregulation, coordinating input from diverse brainstem regions and engaging multiple thermo-effectors to elevate T_c_ (Fig. S7). First, RPa-*vGluT3* neurons can induce piloerection (Fig. 5E), a critical sympathetic heat-retention response^51, 52^, in addition to their previously suggested regulatory roles in BAT thermogenesis, cutaneous vasoconstriction, and lipolysis in white adipose tissue^31, 33, 53–55^. Therefore, RPa-*vGluT3* neurons appear to orchestrate multiple systemic thermogenic responses. Second, extending a previous study implicating the role of RPa neurons in shivering^32^, our study identified a pathway from RPa-*vGluT3* neurons to cIRT, a brainstem structure containing somatic premotor neurons^47, 56, 57^ that contributes to shivering generation (Figs. 5 and 6). Shivering involves rapid alternations between muscle fiber contraction and relaxation, a process that likely requires a premotor pattern-generating network^58, 59^; however, the precise circuit mechanisms remain to be elucidated. As RPa-*vGluT3* neurons also project to other premotor nuclei, such as the rostral IRT and Su5 (Fig.4), these brain regions are implicated in the regulation of the masseter^60, 61^ and facial muscles^62^. These premotor nuclei also send descending projections to spinal motoneurons^56^, suggesting that RPa-*vGluT3* neurons may contribute to shivering in multiple skeletal muscle groups by engaging distinct brainstem premotor circuits.

How do RPa-*vGluT3* neurons regulate diverse thermogenic and heat-retaining mechanisms? Regarding the sympathetic pathways, both piloerection and BAT activation are regulated by postganglionic neurons in the upper thoracic sympathetic trunk^51, 63, 64^. One possibility is that RPa-*vGluT3* neurons regulate a shared sympathetic outflow that simultaneously drives piloerection and the activation of BAT. Alternatively, distinct subpopulations of RPa-*vGluT3* neurons may regulate these two pathways independently. Identifying the precise neuronal subtypes responsible for piloerection and BAT activation is essential to resolving this question. Similarly, it is important to ask whether there is a specific subpopulation of RPa-*vGluT3* neurons responsible for shivering or whether common RPa-*vGluT3* neurons send bifurcated axonal collaterals projecting to both the premotor areas and the spinal cord. Future studies utilizing projection target-initiated axonal mapping^65^ may provide deeper insights into the circuit logic underlying the diverse functions of RPa-*vGluT3* neurons.

### Limitations

While our *vGluT3-Flpo* mice specifically labeled *vGluT3*+ neurons in the RPa, their targeting efficiency was relatively low (approximately 40%; Fig. 1C). This limited efficiency might account for the relatively small effect sizes observed in our experiments. For example, microinjection of bicuculline into the RPa increased T_BAT_ by approximately 3°C in rats^31^, whereas optogenetic or chemogenetic activation of RPa-*vGluT3* neurons in our study led to a 1–2°C increase (Fig. 5D, Fig. S5), although this increase was consistent with a previous study utilizing *vGluT3-Cre* mice^34^. Similarly, chemogenetic inactivation of RPa-*vGluT3* neurons only decreased T_c_ by approximately 1°C (Fig. 1L) and delayed recovery from QIH by approximately 15 min (Fig. 2M), eventually returning to a normal range. This moderate phenotype may be attributed to the limited targeting efficiency of *vGluT3-Flpo* mice, likely due to limited Flpo activity from the bicistronic expression system (Fig. S1). Future studies with a more comprehensive targeting of RPa-*vGluT3* neurons could clarify their relative contributions to thermoregulation. However, we cannot rule out the involvement of additional thermogenic pathways, such as those originating from the paraventricular hypothalamus, in directly targeting sympathetic neurons^66, 67^. In addition, our *Flpo* mice did not specifically target serotonergic *vGluT3*+ Tph2+ neurons in the RPa, reflecting the low expression levels of *vGluT3* in this population. While most serotonergic neurons are distinct from the *vGluT3*+ population in the RPa^68–70^, some have been implicated in BAT-mediated thermogenesis^63, 71^. Thus, their potential contributions to thermogenesis and interaction with *vGluT3*+ Tph2− population warrant further investigation. Furthermore, our circuit mapping excluded local connectivity because of technical limitations (Fig. S4), highlighting the need for future studies to elucidate functional interactions among diverse neuronal subtypes within the RPa. The imaging and manipulation tools established in this study are expected to facilitate molecular- and circuit-level investigations of medullary systems involved in thermoregulation and other homeostatic processes under various conditions, including aging and pathological states.

## Methods

### Key resources table

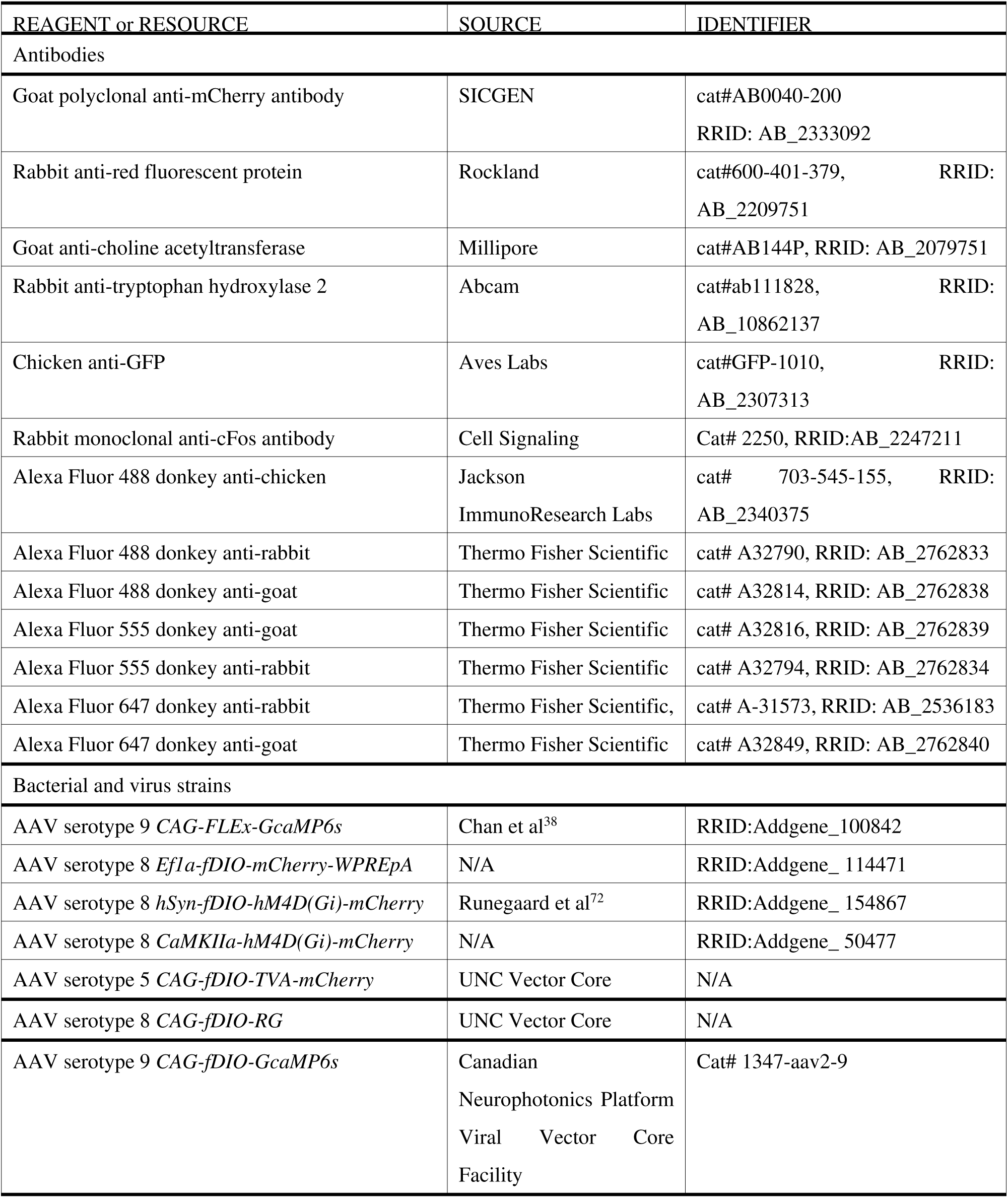
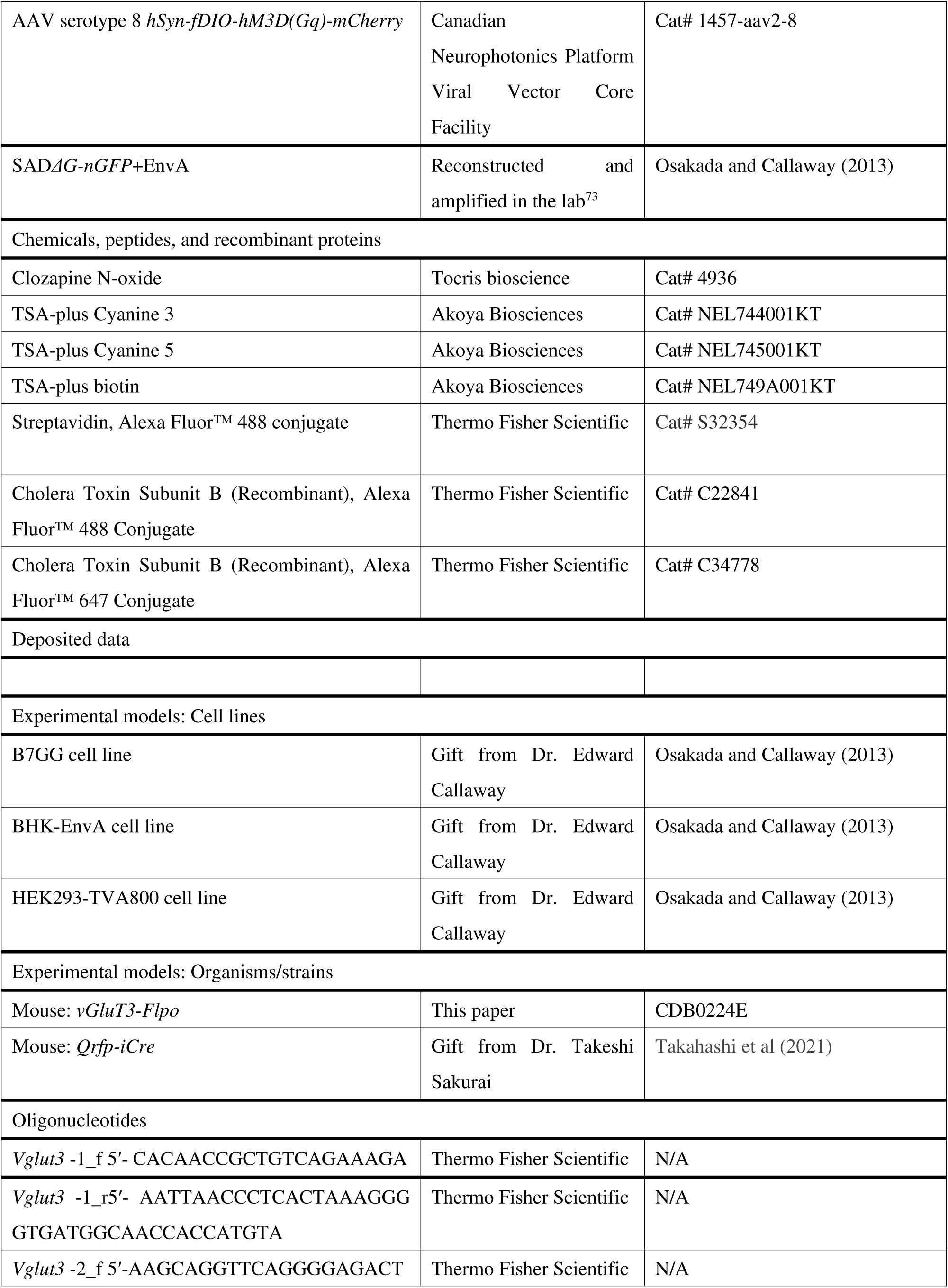
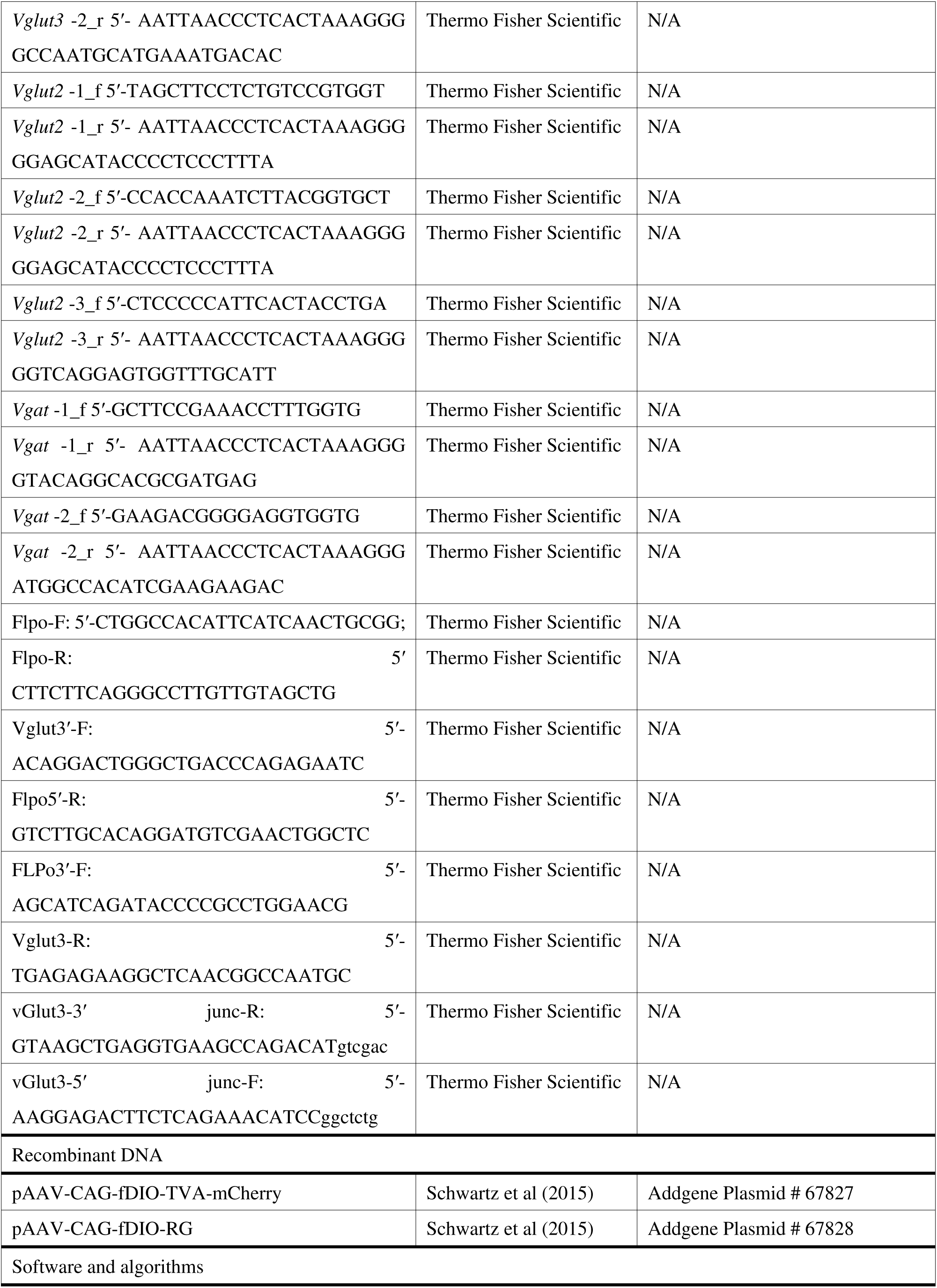
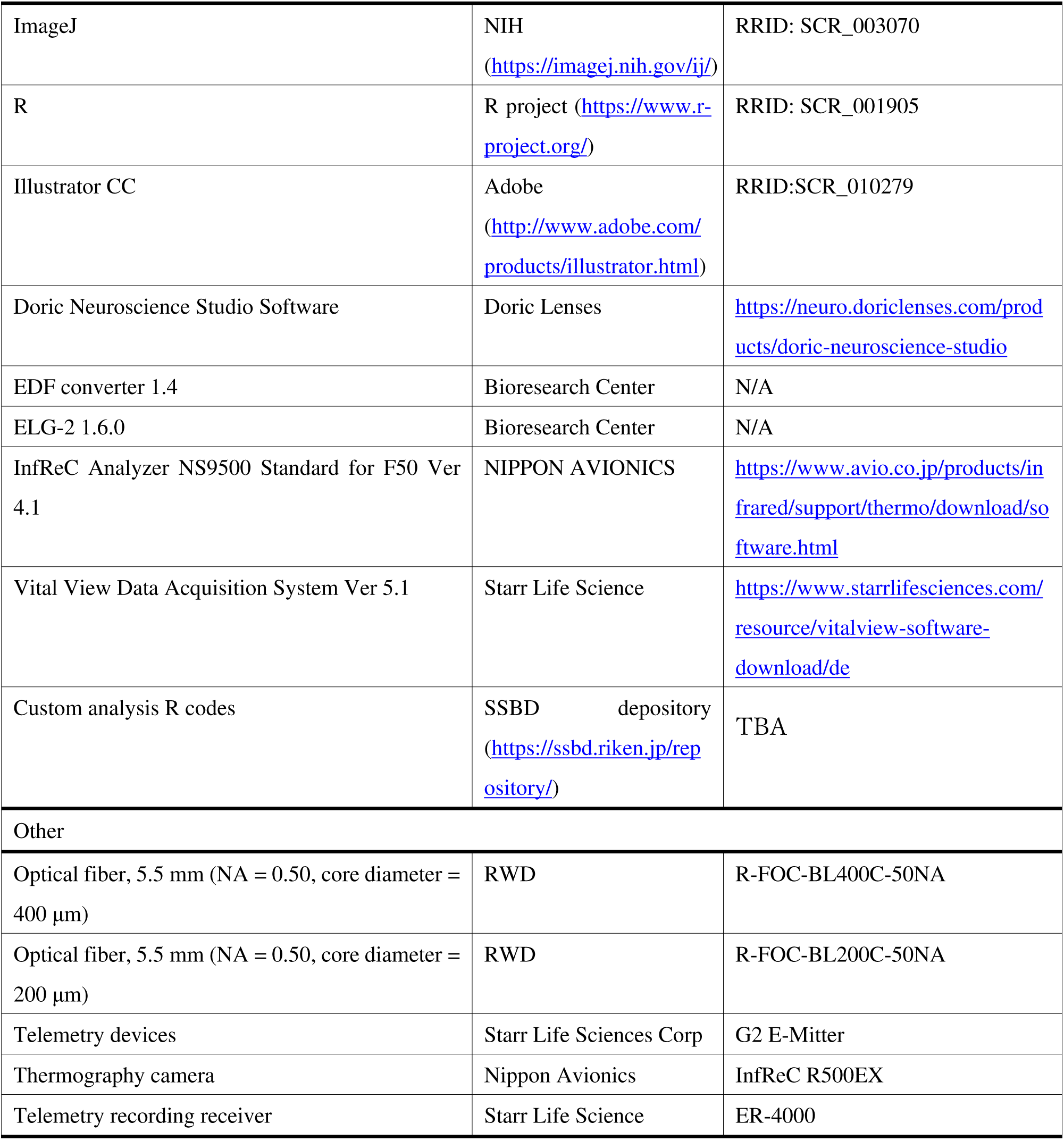

### Animals

All animal experiments were approved by the Institutional Animal Care and Use Committee of the RIKEN Kobe Branch. *Qrfp-iCre* mice have been described previously^16^. *vGluT3-Flpo* mice were originally generated in this study (detailed below). Animals were housed at the animal facility of the RIKEN Center for Biosystems Dynamics Research (BDR) and were fed ad libitum under a 12-h light–dark cycle at an ambient temperature of 20–22°C and humidity levels of 43–57%.

### Generation of *vGluT3-Flpo* knock-in mice

A *vGluT3-Flpo* knock-in mouse line (accession no. CDB0224E, listed at https://large.riken.jp/distribution/mutant-list.html) was generated using CRISPR/Cas9-mediated knock-in techniques in zygotes, as previously described^37^. A donor vector containing *T2A-Flpo* was inserted immediately before the stop codon of *Vglut3* exon 10. The SV40 *nuclear localization signal* (*NLS*) was added to the 5′ end of the *Flpo* open reading frame by polymerase chain reaction (PCR) primers 5′- GGCGCGCCACCATGGCTCCTAAGAAGAAGAGGAAGGTGATGAGCCAGTTCGACATCCTG ; 5′-GTCGACTCAGATCCGCCTGTTGATGTAG. The plasmid *pTCAV-FLEx(loxP)FlpO* (#67829; Addgene) was used as the PCR template. To construct a microhomology-mediated end-joining (MMEJ)-based donor vector, Flpo was cloned into a plasmid harboring a synthetic T2A (*Thoseaasigna* virus 2A) sequence^74^, homology arms, and guide RNA (gRNA) sites (PITCh crRNA3). The gRNA sites were designed using CRISPRdirect^75^ to target regions upstream and downstream of the stop codon (Fig. S1A). For microinjection, a mixture of two CRISPR RNAs (crRNAs) (50 ng/µL), trans-activating crRNA (tracrRNA) (200 ng/µL), donor vector (10 ng/µL), and Cas9 protein (100 ng/µL) was injected into the pronucleus of a C57BL/6 one-cell stage zygote. *Vglut3* crRNA (5′- TCAGAAACATCCTAAATGTCguuuuagagcuaugcuguuuug), PITCh crRNA3 (5′- GCAUCGUACGCGUACGUGUUguuuuagagcuaugcuguuuug), and tracrRNA (5′-AAACAGCAU AGCAAGUUAAAAUAAGGCUAGUCCGUUAUCAACUUGAAAAAGUGGCACCGAGUCGG UGCU) were purchased from FASMAC (Atsugi, Japan). Therefore, 46 F0 founder mice were obtained. Of these, 13 (seven males and six females) were identified as *Flpo*-positive by PCR (Fig. S1B). Further analysis of the 5′ and 3′ junctions and full-length sequencing were conducted on three males and one female (Fig. S1C). The following primers were used for PCR and sequencing analyses.

For the detection of *Flpo* internal sequence:

Flpo-F: 5′-CTGGCCACATTCATCAACTGCGG; Flpo-R: 5′ CTTCTTCAGGGCCTTGTTGTAGCTG.

For the 5′ junction: Vglut3-F: 5′- ACAGGACTGGGCTGACCCAGAGAATC; Flpo5′-R: 5′- GTCTTGCACAGGATGTCGAACTGGCTC.

For the 3′ junction: FLPo3′-F: 5′- AGCATCAGATACCCCGCCTGGAACG; Vglut3-R: 5′- TGAGAGAAGGCTCAACGGCCAATGC.

For the sequence analysis, the PCR products were obtained using the primers vGlut3-F and vGlut3-3′ junc-R: 5′- GTAAGCTGAGGTGAAGCCAGACATgtcgac or vGlut3-5′ junc-F: 5′- AAGGAGACTTCTCAGAAACATCCggctctg and vGlut3-R. The PCR products were subcloned into the pCR Blunt II TOPO vector (Zero Blunt TOPO PCR Cloning Kit, Thermo Fisher Scientific) and sequenced using the M13-Forward and M13-Reverse primers (Fig. S1D).

Germline transmission of the *vGlut3-Flpo* allele was confirmed by genotyping the F1 mice (Fig. S1E). The line was established using a male that harbored the target sequence, as verified by PCR and sequencing. Genotyping PCR was performed using Flpo-F and Flpo-R internal primers, as described above.

### Viral preparations

AAV vectors were purchased from Addgene, with titers represented as genome particles (gp) per mL: AAV serotype 8 *Ef1a-fDIO-mCherry-WPREpA* (1.8 × 10^13^ gp/mL, #114471-AAV8)

AAV serotype 8 *hSyn-fDIO-hM4D(Gi)-mCherry* (5.0 × 10^12^ gp/mL, #154867-AAV8). AAV serotype 8 *CaMKIIa-hM4D(Gi)-mCherry* (2.4 × 10^13^ gp/mL, # 50477-AAV8).

AAV vectors were generated by the University of North Carolina Vector Core, using plasmids purchased from Addgene.

AAV serotype 5 *CAG-fDIO-TVA-mCherry* (2.4 × 10^13^ gp/mL, # 67827)

AAV serotype 8 *CAG-fDIO-RG* (1.0 × 10^12^ gp/mL, #67828).

AAV vectors were purchased from the Canadian Neurophotonics Platform Viral Vector Core Facility (RRID: SCR_016477).

AAV serotype 9 *CAG-fDIO-GcaMP6s* (6.9 × 10^12^ gp/mL, construct-1347-aav2-9);

AAV serotype 8 *hSyn-fDIO-hM3D(Gq)-mCherry* (4.7 × 10^12^ gp/mL, construct-1457-aav2-8).

RV*ΔG-nGFP* preparation was carried out using cell lines as previously described^73^. The histone-GFP fragment was amplified from the *pAAV-TRE-HTG* plasmid (Addgene # 27437) using the following primers:

H2B-arm-F: 5′- atccctcaaaggacctgcaggGCCACCATGCCAGAGCCAG;

EGFP-arm-R: 5′- gactgaaaagctaccgcggTTACTTGTACAGCTCGTCCATGCCGA.

The PCR product was subcloned into the *pSAD-dG-F3* vector (a gift from Ed Callaway)^73^, which was treated with SbfI and SacII restriction enzymes using the In-Fusion HD Cloning Kit (639648, Takara) to yield the *pSAD-dG-F3-nGFP* plasmid. RV*Δ*G-nGFP was generated de novo using the B7GG cell line (a gift from Ed Callaway) along with *pCAG-B19N*, *pCAG-B19P*, *pCAG-B19L*, *pCAG-B19G* (gifts from Ed Callaway), and *pSAD-dG-F3-nGFP*. Viral particles were pseudotyped using BHK-EnvA cells (a gift from Ed Callaway). The titer of RV*Δ*G-nGFP+EnvA used in this study was estimated to be 3.7×10^9^ infectious particles per mL, based on serial dilutions of the virus stock followed by infection of the HEK293-TVA800 cell line (a gift from Ed Callaway).

### Virus injection

Mice were deeply anesthetized by intraperitoneal injection of 65 mg/kg ketamine (4987081519033, Daiichi Sankyo) and 13 mg/kg xylazine (X1251, Sigma-Aldrich) and then positioned in a stereotactic frame (cat#68045, RWD). The skull was positioned such that the dorsal surface was level with the heights of the Bregma and Lambda aligned. For cIRT injections, the skull was angled such that Lambda was positioned 0.6 mm lower than Bregma. A glass micropipette filled with AAV was then placed into the target region using the following coordinates: AVPe, 0.8 mm anterior, 0.25 mm lateral, and 5.5 mm ventral from the Bregma; RPa, 6.2 mm posterior, 0.0 mm lateral from the Bregma, and 5.3 mm ventral from the brain surface; cIRT, 6.4 mm posterior and 1.2 mm lateral from the Bregma, and 5.3 mm ventral from the brain surface. A 240 nl solution of AAV was injected at a speed of 80 nl/min using a micropump (World Precision Instruments, UMP3T-1). After the viral injection, the animals were returned to their home cages.

In Fig. 4, a cocktail of AAV5-*CAG-fDIO-TVA-mCherry* and AAV8-*CAG-fDIO-RG* (240 nl, mixed at a 1:4 ratio) was injected into the RPa of *vGluT3-Flpo* mice. Three weeks after the injection, 240 nl of SAD*ΔG-nGFP*+EnvA was injected into the same region. The mice were kept in their home cages for 7 days before perfusion. For the control experiment (Fig. S4), AAV8*-CAG-fDIO-RG* was omitted to assess the extent of glycoprotein-independent nonspecific labeling of the RV.

### Injection of cholera toxin subunit B

To retrogradely label iBAT-projecting and piloerector muscle-projecting sympathetic postganglionic neurons (Fig. S6A and S6B), cholera toxin subunit B (CTB) was injected into the target organs of *vGluT3-Flpo* mice following optogenetic activation experiments (Fig. 5). CTB Alexa Fluor™ 488 Conjugate (CTB488; C22841, Thermo Fisher Scientific) or CTB Alexa Fluor™ 647 Conjugate (CTB647; C34778, Thermo Fisher Scientific) was prepared as a 0.1% solution in phosphate buffered saline (PBS).

Throughout the procedure, the animals were maintained under anesthesia using intraperitoneal injections of ketamine and xylazine, as described in the virus injection section. For iBAT tracing, the iBAT was exposed via an intrascapular incision, and 4 μl of CTB647 (1 μl per injection, two injections per side) was delivered using a pulled glass pipette. For piloerector muscle tracing, 6 μl of CTB488 (1 μl per injection, six injections in total) was injected intradermally. The injection sites were distributed along the dorsal midline of the cervical skin, spanning spinal levels C4 to Th2, with approximately 2 mm spacing between adjacent sites. After surgery, the mice were single-housed and sacrificed six days later.

### Fiber implantation

At least two days after virus injection, the mice were anesthetized and placed in a stereotactic apparatus, as mentioned above. An optical fiber (R-FOC-BL400C-50NA, RWD) for imaging experiments or an optic cannula (R-FOC-BL200C-50NA, RWD) for optogenetic manipulation was placed above the RPa (midline, 6.2 mm posterior and 6.3 mm ventral from the Bregma) or AVPe (midline, 0.8 mm anterior and 5.2 mm ventral from the Bregma) and affixed to the skull with dental cement.

### Cold challenge

To examine the photometric signals of RPa-*vGluT3* neurons when mice were exposed to a cold environment, the cooled plates (0°C; 188 mm × 98 mm × 24 mm, DAISO) were placed beneath the cage for 25 min. This reduced the temperature of the cage floor to 15°C within 15 min. The last 10 min of exposure were analyzed. For control experiments, identical plates were maintained at room temperature.

### Body temperature and locomotor activity monitoring

To monitor T_c_ and locomotor activity, telemetry devices (G2 E-Mitter, Starr Life Sciences Corp.) were surgically implanted in mice following a two-week recovery period after AAV injection or fiber implantation.

For surgical implantation, mice were anesthetized by intraperitoneal injection of 65 mg/kg ketamine and 13 mg/kg xylazine. A medial incision was made in the abdominal cavity, and a telemetry temperature transmitter was placed inside. After filling the abdominal cavity with saline (Ootsuka), the peritoneum and abdominal skin were sutured with coated Vicryl sutures (D5893; Johnson & Johnson). Gentamicin (5 mg/kg; Takata Pharmaceutical) was administered intraperitoneally immediately after surgery. Mice were kept in their home cages for at least 5 days before recording.

The day before recording, the mice and their home cages were placed on a recording receiver (ER-4000, Starr Life Science). For imaging or optogenetic experiments, the mice were transferred to acrylic cages (27 cm × 19 cm × 25 cm; Sanplatic, Japan) and positioned on the recording receiver. Body temperature and locomotor activity were recorded every minute using recording software (Vital View, Starr Life Science).

To monitor T_BAT_, the mice were monitored using a thermography camera (InfReC R500EX, Nippon Avionics) placed 20 cm above the cage floor. The back hair of each mouse was shaved to facilitate T_BAT_ detection. Thermograms were recorded every minute and the maximum temperature in each frame was used as the T_BAT_ of the mouse. Mice with T_BAT_ values below 34°C were excluded from analysis. In Fig. S3, T_BAT_ data from one mouse were unavailable because of a recording failure caused by battery depletion, whereas the T_c_ data were successfully recorded. Consequently, the number of animals included in the analyses differed between the T_c_ and T_BAT_.

### Section preparation

Mice were deeply anesthetized with isoflurane and perfused with 10 mL PBS, followed by 50 mL 4% PFA. The brains were extracted, and the thoracic spines were removed from the mediastinal organs and adipose tissues. The specimens were post-fixed overnight at 4°C in 4% PFA and then immersed in a 30% sucrose solution in PBS at 4°C until they sank. The spinal cord was dissected from the vertebrae using micro-scissors. The stellate ganglion attached to the vertebrae was decalcified by immersion in 0.5 M ethylenediaminetetraacetic acid (EDTA) (pH 8.0) overnight at 4°C, followed by cryoprotection in 30% sucrose. The level of the thoracic spinal cord was determined based on the vertebrae position, as previously reported^76^. The specimens were embedded in optimal cutting temperature compound (#4583, Tissue-Tek) and sectioned serially at 60 µm for immunohistochemistry or 30 µm for *in situ* hybridization using a cryostat (Leica, Germany).

### Immunohistochemistry

Serial brain and spinal cord sections were collected in 24-well plates, and every third section was processed to immunohistochemistry. Sections were washed three times with PBS, then immersed in 1% Triton in PBS at room temperature for 3 h. The sections were blocked with 10% Blocking One (cat#03953-95, Nacalai Tesque) in PBS containing 0.3% Triton X-100 (blocking solution) at room temperature for 1 h. They were then incubated with the primary antibody in a blocking solution at 4°C overnight. Following this, sections were rinsed three times with PBS and incubated with the secondary antibody at 4°C overnight. Finally, the sections were rinsed with PBS, mounted, and cover slipped using Fluoromount (cat#K024; Diagnostic BioSystems). The primary antibodies used in this study were: Goat anti-mCherry (1:1000; cat#AB0040-200, SICGEN, RRID: AB_2333092); Rabbit anti-red fluorescent protein (1:1000; cat#600-401-379, Rockland, RRID: AB_2209751); Goat anti-choline acetyltransferase (1:500; cat#AB144P, Millipore, RRID: AB_2079751); Rabbit anti-tryptophan hydroxylase 2 (1:500; cat#ab111828, Abcam, RRID: AB_10862137); Chicken anti-GFP (1:2000, cat#GFP-1010, Aves Labs, RRID: AB_2307313); and Rabbit anti-cFos (1:2000; cat#2250, Cell Signaling, RRID:AB_2247211). The secondary antibodies were: Alexa Fluor 488 donkey anti-chicken (1:500; cat# 703-545-155, Jackson ImmunoResearch Labs, RRID: AB_2340375); Alexa Fluor 488 donkey anti-rabbit (1:500; cat# A32790, Thermo Fisher Scientific, RRID: AB_2762833); Alexa Fluor 488 donkey anti-goat (1:500; cat# A32814, Thermo Fisher Scientific, RRID: AB_2762838); Alexa Fluor 555 donkey anti-goat (1:500; cat# A32816, Thermo Fisher Scientific, RRID: AB_2762839); Alexa Fluor 555 donkey anti-rabbit (1:500; cat# A32794, Thermo Fisher Scientific, RRID: AB_2762834); Alexa Fluor 647 donkey anti-rabbit (1:500; cat# A-31573, Thermo Fisher Scientific, RRID: AB_2536183); and Alexa Fluor 647 donkey anti-goat (1:500; cat# A32849, Thermo Fisher Scientific, RRID: AB_2762840).

Representative images were obtained using a confocal microscope (Zeiss Axioscan7 or Leica SP8).

### *In situ* hybridization

To generate cRNA probes, DNA templates were amplified from spinal cord cDNA using PCR (cat#MD-23; Genostaff). A T3 RNA polymerase recognition site (5′- AATTAACCCTCACTAAAGGG) was added to the 3′ end of the reverse primers. The primer sets and sequences of the probe targets were as follows.

*Vglut3* -1: 5′-CACAACCGCTGTCAGAAAGA; 5′-GTGATGGCAACCACCATGTA

*Vglut3* -2: 5′-AAGCAGGTTCAGGGGAGACT; 5′-GCCAATGCATGAAATGACAC

*Vglut2* -1: 5′-TAGCTTCCTCTGTCCGTGGT; 5′-GGGCCAAAATCCTTTGTTTT

*Vglut2* -2: 5′-CCACCAAATCTTACGGTGCT; 5′-GGAGCATACCCCTCCCTTTA

*Vglut2* -3: 5′-CTCCCCCATTCACTACCTGA; 5′- GGTCAGGAGTGGTTTGCATT

*Vgat* -1: 5′-GCTTCCGAAACCTTTGGTG; 5′-GTACAGGCACGCGATGAG

*Vgat* -2: 5′-GAAGACGGGGAGGTGGTG; 5′-ATGGCCACATCGAAGAAGAC

*In vitro* transcription reactions were performed according to the manufacturer’s instructions (Roche Applied Sciences). First, 600–1000 ng of DNA for these genes was incubated with Dig (Digoxigenin) (cat#11277073910)- or fluorescein (cat#11685619910)-RNA labeling mix, T3 RNA polymerase (cat#11031163001), and Rnase inhibitor (cat#3335399001) at 37°C for 6 h. After incubation with Dnase I (Promega, cat#M6101) for another 20 min at 37°C, followed by EDTA treatment (cat#AM9260G, Life Technologies), cRNA probes were purified using ProbeQuant G-50 Micro Columns (cat#28-9034-08, Cytiva).

Fluorescent *in situ* hybridization (ISH) combined with anti-GFP and anti-TPH2 immunohistochemical staining was performed as previously reported^77^. For detecting *Vglut3* mRNA (Fig. 1B–1D), after hybridization and washing, brain sections were incubated with horseradish peroxidase (HRP)-conjugated anti-Dig (1:500; cat#1120773390, Roche Applied Science) antibody overnight at 4 °C. The following day, the signals were amplified with TSA-plus Cyanine 3 (1:70 in 1x plus amplification diluent; cat#NEL744001KT; Akoya Biosciences) for 25 min. After washing with PBS containing 0.1% Tween-20 (PBST) for 5 min, the sections were incubated with anti-GFP (1:2000; cat#GFP-1010, Aves Labs) and anti-TPH2 (1:500; cat#ab111828, Abcam, RRID: AB_10862137) at 4°C overnight. GFP-positive and TPH2-positive cells were visualized using anti-chicken Alexa Fluor 488 (cat#703-545-155, Jackson ImmunoResearch, 1:250) and Alexa Fluor 647 donkey anti-rabbit antibodies (1:500; cat# A-31573, Thermo Fisher Scientific, RRID: AB_2536183).

For dual-color ISH combined with anti-GFP staining (Fig. 3I), an HRP-conjugated anti-flu antibody (1:250; cat#NEF710001EA, Akoya Biosciences) was used to detect Flu-labeled RNA probes using TSA-plus Cyanine 3 (1:70 in 1× plus amplification diluent; cat#NEL744001KT, Akoya Biosciences) for 25 min. After a 5-min PBST wash, HRP was inactivated with a 2% sodium azide solution in PBS for 15 min at room temperature, followed by five 5-min washes with PBST. The sections were then incubated with HRP-conjugated anti-Dig (1:500) and anti-GFP (cat#GFP-1010, Aves Labs, 1:1000) antibodies at 4°C overnight. Signals were amplified using TSA-plus Cyanine 5 (cat#NEL744001KT, Akoya Biosciences; 1:70 in 1× plus amplification diluent) for 25 min, followed by washing. GFP-positive cells were visualized using anti-chicken Alexa Fluor 488 antibody (1:250, cat#703-545-155, Jackson ImmunoResearch). Nuclei were counterstained with PBS containing 50 ng/mL 4’, 6-diamidino-2-phenylindole dihydrochloride (DAPI; cat #D8417, Sigma-Aldrich). Images were acquired using an Olympus BX53 microscope equipped with a 10x (N.A. 0.4) objective lens. The cells were then counted manually.

### Optogenetic stimulation and chemogenetic manipulations

The fiberoptic cannulas implanted in the mice were connected to a fiberoptic patch cable (200 µm diameter, NA: 0.22, 1.0 m length, Doric Lenses) using ceramic sleeves (Thorlabs). The mice were allowed to habituate for at least 1 h before stimulation. Diode-pumped solid-state lasers (465 nm blue, IOS-465, RWD) were used for optogenetic manipulation. The laser output at the optical fiber tip was measured using a laser checker (PM100D, Thorlabs) and was adjusted to 6–8 mW. Laser stimulation was applied at 2 Hz with a 10-ms pulse width for 3 h in the QIH experiment (Fig. 2) and at 40 Hz with a 10-ms pulse width for 10 min to stimulate the RPa-*vGluT3* neurons (Figs. 5 and 6). In Fig. 1I–1M, mice received either saline or CNO (2 mg/kg, #4936, Tocris Bioscience) and their T_c_ was monitored at room temperature or 4°C as indicated in the figure panel. As shown in Fig. 2, the mice received either saline or CNO 30 min after the induction of QIH by laser stimulation on day 1, with treatment conditions counterbalanced on day 2. One mouse showing the spontaneous T_c_ recovery and two mice that failed to reach a minimum T_c_ below 27°C were excluded from the analysis. Rewarming speed was calculated from the time of the laser termination to the point at which T_c_ reached 35°C. As shown in Fig. 6, the mice were injected with either saline or CNO (2 mg/kg) 45–60 min after optogenetic stimulation on day 1, with the treatment conditions counterbalanced on day 2. The CNO was dissolved in saline at a concentration of 0.5 mg/mL.

### Fiber photometry recording

Fiber photometry recordings were performed by delivering excitation lights (465 nm modulated at 309.944 Hz and 405 nm modulated at 208.616 Hz) and collecting the emitted fluorescence using an integrated fluorescence mini-cube (Doric, iFMC4_AE(405)_E(460–490)_F(500–550)_S). Light collection and demodulation were performed using a Doric Photometry Setup and Doric Neuroscience Studio Software (Doric Lenses). The 405 nm signal was recorded as an isosbestic signal (non–calcium-dependent), and the 465-nm signal was recorded as a calcium-dependent GCaMP6s signal. The power output at the fiber tip was approximately 5–10 µW. The signals were initially acquired at 12 kHz and then decimated to 120 Hz for recording. All optical components were purchased from Doric Lenses (Quebec, Canada).

### Analysis of Ca^2+^ imaging data

Data processing was performed using custom-made R-code following the methodologies outlined in previous study^78, 79^. Both the 405-nm and 465-nm signals were subjected to low-pass filtering at 5 Hz and high-pass filtering at 0.001 Hz. In the QIH Ca^2+^ recording experiment (Fig. 2), both signals were high-pass filtered at 0.005 Hz owing to temperature-induced baseline fluctuations in the fluorescent signals^80^. To remove motion noise, a control signal was calculated by fitting the Ca^2+^-independent 405-nm signal to the Ca^2+^-dependent 465-nm signal using least squares linear regression. This fitted control signal was subtracted from the 465-nm signals, and the resulting residual was normalized to the Z-score.

An “upregulated signal” was defined as a peak that surpassed 3 SD and was sustained above 0 SD for at least 3 s. “Up” or “Down” phases were defined as periods during which temperature continuously rose or fell ≥5 min, with a minimum temperature change of 0.3°C. The pre-up phase was defined as the 4 min preceding the up phase; if it overlapped with the up phase, it was considered part of the up phase. The AUC for upregulated signals during each phase was measured. Mice exhibiting upregulated signals lasting <3 min/h were excluded from the analysis. Time zero in the T_c_ analysis following the termination of QIH (Fig. 2G) was defined as the moment when the regression line of increasing T_c_ intersected the baseline T_c_ during QIH.

### Counting input neurons to RPa-*vGluT3* neurons

Images of every third 60 µm whole-brain sections were obtained by slide scanner (Zess Axioscan7, Zeiss) after immunostaining. The number of starter (GFP and mCherry double-positive) and input neurons (GFP-positive) were counted manually and assigned to brain areas based on the classification of the Allen Mouse Brain Atlas. The convergence index was calculated by dividing the number of input neurons in each brain area by the total number of starter neurons.

### Axon density analysis

Every third 60 µm coronal section of the whole brain, as well as spinal cord sections spanning from the thoracic segment 1 (Th1) to the lumbar segment 1 (L1), was imaged using a slide scanner (Zeiss Axioscan 7) after immunostaining. To quantify the density of mCherry-positive axons originating from RPa-*vGluT3* neurons (Fig. 4), regions of interest (ROIs) were manually delineated for the relevant brain and spinal cord regions. The mCherry channel in each image was binarized using the Renyi entropy thresholding plug-in function in the ImageJ (Fiji). For each mouse, the mCherry-positive area within each ROI was normalized to the region with the maximum signal to calculate relative axon density. Although mCherry-positive axons were detected in white matter tracts such as the ventral spinocerebellar tract in the brainstem and the lateral funiculus in the spinal cord, these areas were excluded from the analysis because they likely represent passing fibers rather than terminal projections.

### EMG recording

A handmade EMG electrode was implanted at the same time as fiber implantation. Three silver wires including a ground electrode were placed under the nuchal muscles and fixed with dental cement. On the recording day, the mouse with the implanted EMG electrode was connected to a recorder (ELG-2, Bioresearch Center) and habituated for at least 1 h before laser stimulation.

To improve the signal-to-noise ratio, EMG was performed under anesthesia. The mice were initially anesthetized with 3% isoflurane for 5 min in an anesthesia box (Natsume, Cat#KN-1010-S) and transferred to an anesthesia apparatus (Natsume, Cat#KN-1019-1) containing 0.6% isoflurane. Throughout the recording, mice were placed on a digital hot plate (Corning, PC-420D) to maintain body temperature at 35°C. The optical fiber was connected to a patch cable, and laser stimulation was applied 1 min after 0.6% isoflurane anesthesia.

Data were recorded at a sampling rate of 100 Hz and converted to text format using the EDF-converter software (Bioresearch Center). The recordings were high-pass filtered at 10 Hz and rectified to absolute values using R software. The integral of the EMG amplitude was calculated in 10-s bins, and the EMG values were normalized to the minimum amplitude observed during the 1-min baseline period prior to laser stimulation. Mice exhibiting excessive baseline noise in EMG recordings, defined as average EMG amplitudes exceeding 100 µV even under isoflurane anesthesia, were excluded from the analysis because of poor signal quality.

### Measurement of cervical twitching behavior and piloerection

For the optogenetic activation of RPa*-vGluT3* neurons (Fig. 5), video recordings of the mice were taken for 12 min, including 10 min of laser stimulation and 1 min before and after stimulation. The number of cervical twitching behaviors was manually counted every minute. Cervical twitching was defined as bending of the cervical lordosis (Movie S1), excluding movements related to grooming or eating pellets.

Piloerection was assessed by measuring changes in the angles of the cervicothoracic hair before and after stimulation (Fig. 6E). Specifically, hair snapshots were taken and angle changes were analyzed using the “Find Edges” function in ImageJ (Fiji). For the pre-laser stimulation condition, angle changes were calculated over a baseline window from −45 s to −15 s relative to the onset of laser stimulation. For the laser stimulation condition, changes were measured between −15 s and +15 s. From each mouse, five individual hairs were selected for measurement and subsequent analysis.

### Statistical analysis

The statistical analyses for each experiment, including the specific statistical tests used and the exact number of animals, are detailed in each figure legend. P-values are reported in figure legends or panels; non-significant values are not noted. Normality and homogeneity of variance were assessed using the Shapiro–Wilk normality test and F-test, respectively. For the details of the statistics, please see the Statistical Table.

### Data and materials availability

All fiber photometry data will be deposited in the SSBD repository and will be publicly accessible upon publication. All other data are available in the main manuscript and supplementary material. All materials, including *vGluT3-Flpo* mice, are available from the corresponding authors upon request.

## Author contributions

S.U. and K.M. conceived the experiments. S.U. performed the experiments and analyzed the data with technical support from M.H. *Qrfp-iCre* mice were provided by T.S. *vGluT3-Flpo* mice were generated by T.A. and K.I. with technical support from M.H. S.U. and K.M. wrote the paper with contributions from all co-authors.

## Acknowledgments

We thank the staff at the RIKEN BDR animal facility for animal care and *in vitro* fertilization; Shigefumi Yokota and members of the Miyamichi Laboratory for critically reading the manuscript; Addgene, the University of North Carolina Vector Core, and the Canadian Neurophotonics Platform Viral Vector Core Facility for AAV production; and Satsuki Irie for technical assistance. This study was supported by the RIKEN Special Postdoctoral Researchers Program and JSPS KAKENHI (22K15237, 25K18589) to S. U., and the JST CREST Program (JPMJCR2021) and JSPS Transformative Research Areas (A) (23H04945, 23H04939) to K. M.

## Supplementary Figures

**Figure S1:**
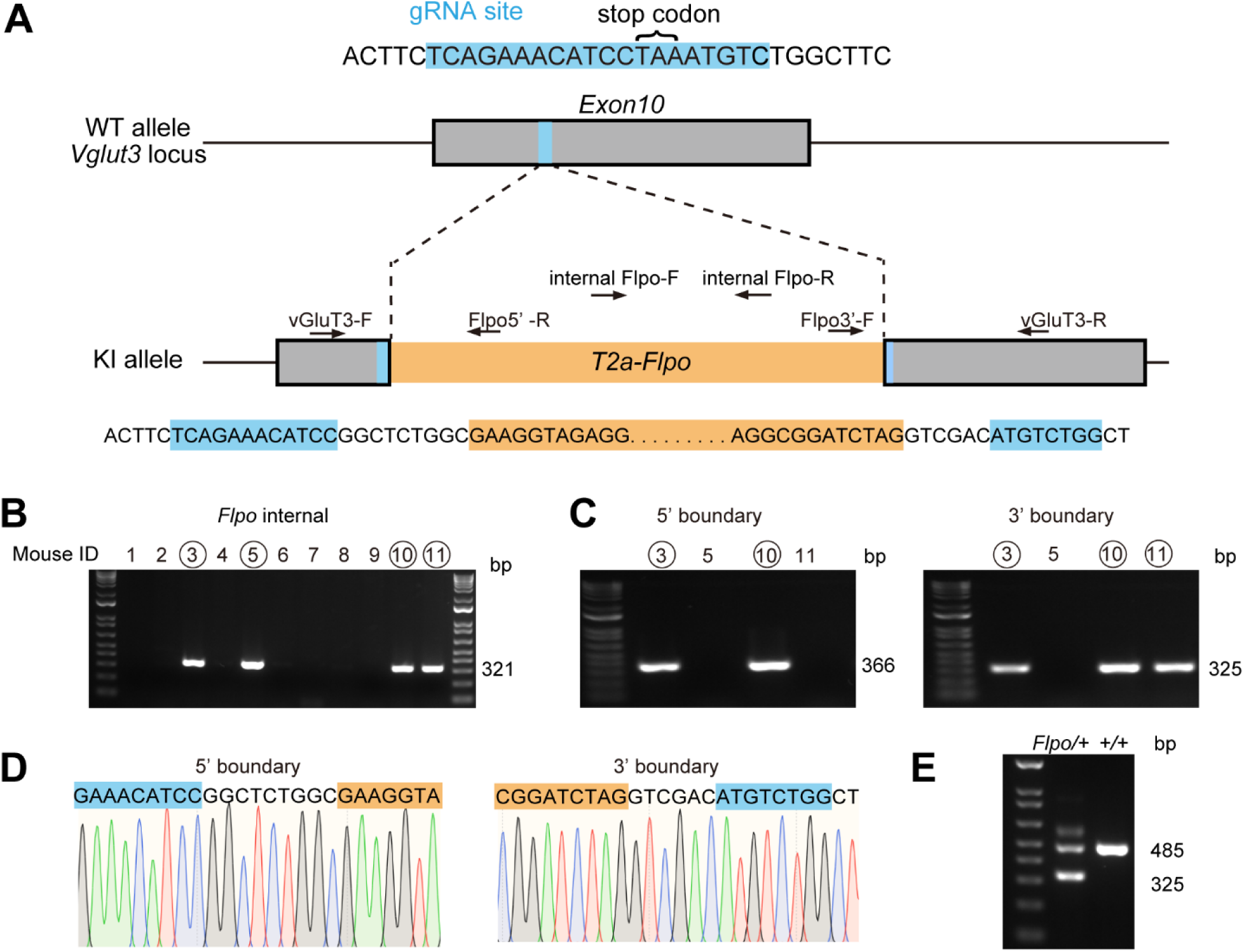
Generation of *vGluT3-Flpo* mice, related to Fig. 1. (A) Schematic of the knock-in (KI) strategy. A *T2A-Flpo* cassette was inserted into the coding end located in exon 10 of the *vGluT3* (*Slc17a8*) locus. Primer locations are indicated by arrows. (B) Screening of *vGluT3-Flpo* founder mice. The initial screening was performed using an internal *Flpo* sequence. The size (base pair, bp) of the PCR product is indicated to the right of the gel images. (C) *Flpo*-positive mice were further screened by amplifying the 5′ and 3′ boundaries. PCR primers used were as follows: *vGluT3-F*: *Flpo-5′R* for the 5′ boundary and *Flpo-3′F*: *vGluT3-R* for the 3′ boundary. (D) Confirmation of the 5′ and 3′ boundary sequences. (E) Representative electrophoresis gel image to examine the KI allele (left) and the WT allele (right). PCR primers *vGluT3-F*: *Flpo-3′F*: *vGluT3-R* were used.

**Figure S2:**
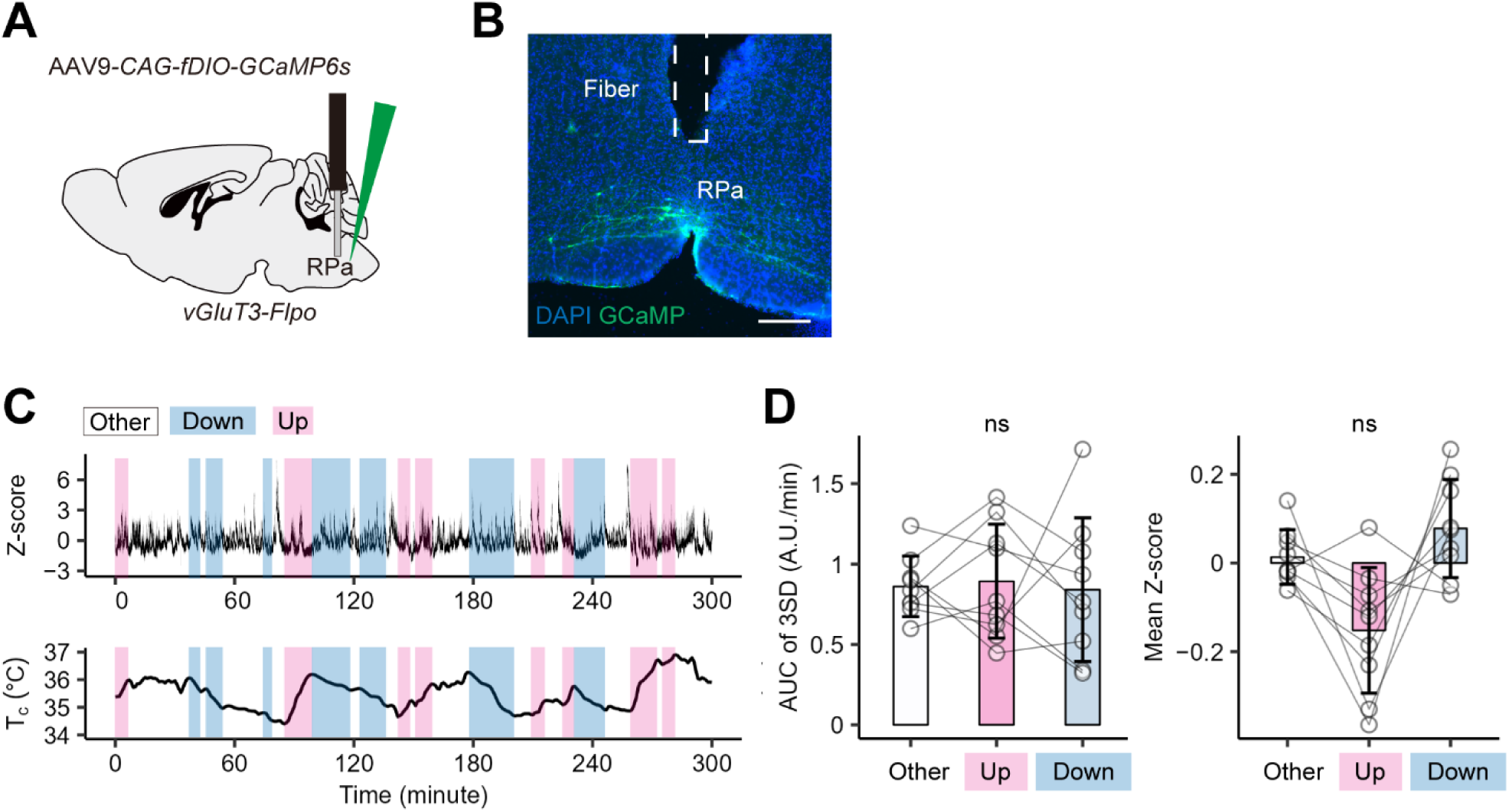
Quantification of photometric signals from RPa-*vGluT3* neurons, related to Fig. 1. (A) Schematic of virus injection. (B) Representative coronal section of the RPa showing GCaMP6s expression (green) and fiber location, with DAPI nuclear staining (blue). Scale bars, 100 µm. (C) Representative 7-h photometry trace (top) and T_c_ trace (bottom). Magenta- and blue-shaded regions represent up and Down phases, respectively. This panel presents the same dataset as shown in Fig. 1D. (D) AUC of 3SD signals (left) and mean Z-score (right). No significant differences were observed among the states, as determined by the Wilcoxon signed-rank test with Bonferroni correction. These data indicate that RPa-*vGluT3* neurons are not active during the Up phase. Error bars indicates SD.

**Figure S3:**
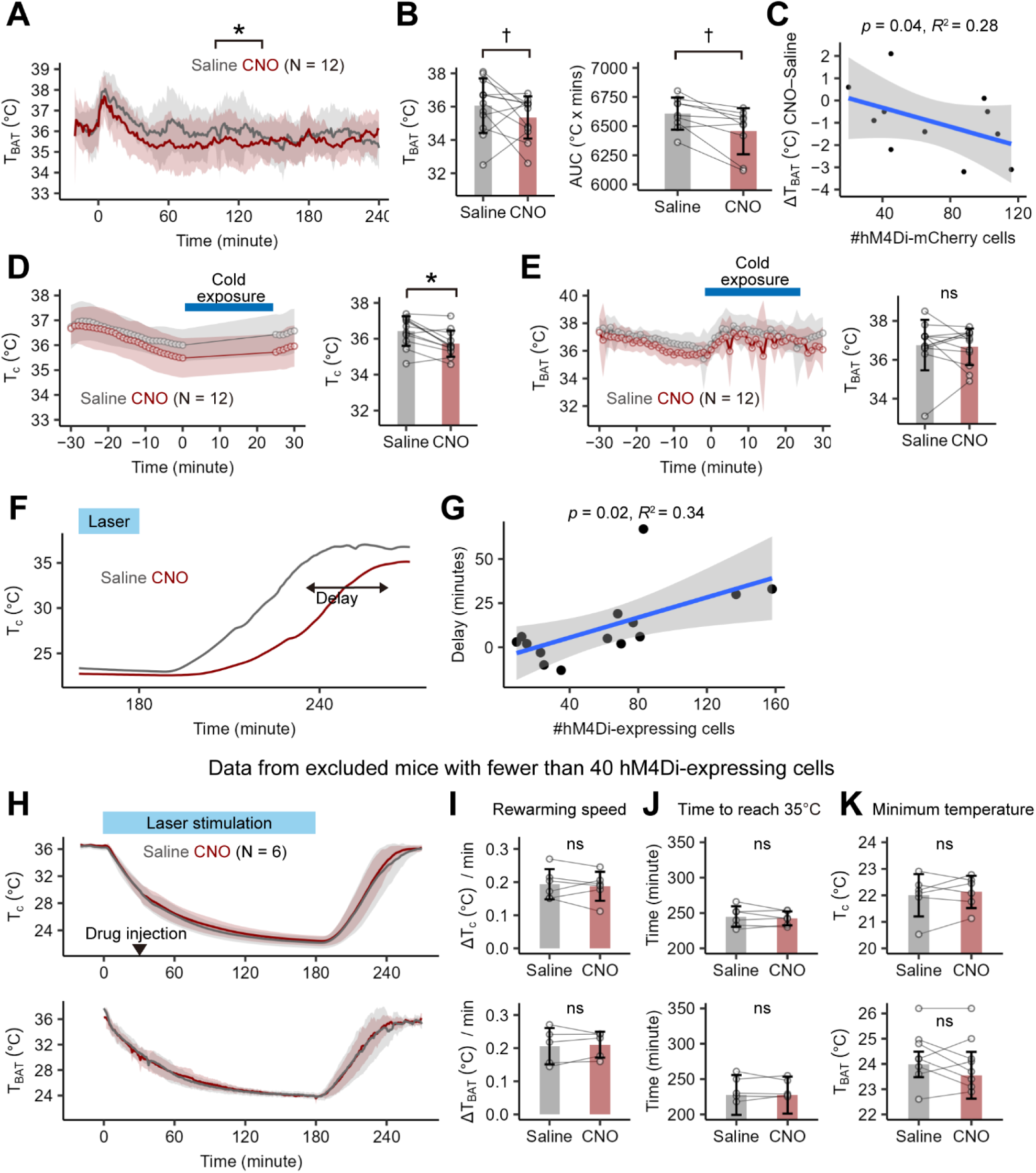
Additional data on chemogenetic inhibition of RPa-*vGluT3* neurons during basal state and QIH, related to Figs. 1 and Fig. 2. (A) Group mean traces of T_BAT_ in hM4Di+ mice following administration of saline (gray) or CNO (red) at time 0. Repeated-measures two-way ANOVA shows significant solution (*p* < 0.01), time course (*p* < 0.01), and interaction (*p* < 0.01) effects. (B) T_BAT_ at 120 min following saline or CNO injection (left) and AUC for T_c_ from 0 to 240 min (right). Unexcluded data showed no significant difference between saline and CNO injections; however, a statistically significant difference emerged when the two mice with fewer than 40 hM4Di-expressing cells were excluded. † *p* < 0.05, Wilcoxon signed-rank test. N = 10. (C) Correlation between the number of hM4Di-mCherry+ cells and the change in T_BAT_ at 120 min following saline or CNO injection. Adjusted coefficient of determination (R^2^) is shown, with the *p*-value calculated using a *t*-test under the null hypothesis of no correlation. (D, E) Group mean traces of T_c_ (D) and T_BAT_ (E) following cold exposure (blue bar). CNO (red) or saline (gray) was administered 30 min before cold exposure. Lines represent group means; shading denotes SD. N = 12. Right panels show T_c_ (D) and T_BAT_ (E) at 25 min following saline or CNO injection. *, *p* < 0.05 by Wilcoxon signed-rank sum test. N = 12. (F) Representative T_c_ curve during recovery from QIH. The delay was defined as the difference in the latency to reach 35°C between saline- and CNO-injected conditions. (G) Correlation between the number of hM4Di-mCherry+ cells and the rewarming delay. The adjusted coefficient of determination (R^2^) is shown, with the p-value calculated using a *t*-test under the null hypothesis of no correlation. (H–K) Same analyses as in Fig. 2K–N, but using animals with ≤ 40 hM4Di-mCherry+ neurons in the RPa. In these mice, CNO administration had no significant effect on the recovery from QIH. N = 6 for T_c_ data, and N = 5 for _BAT_ data. (H) T_c_ (top) and T_BAT_ (bottom) traces after laser stimulation (blue bar). CNO (red) or saline (gray) was administered 2.5 h before laser cessation. Lines represent group means; shading denotes SD. Repeated-measures two-way ANOVA: Drug effect, n.s.; time effect, *p* < 0.01; interaction effect, n.s. (I–K) Rewarming speed (I), latency to reach 35°C (J), and minimum temperature (K) of T_c_ (top) and T_BAT_ (bottom) after saline or CNO injection. ns: non-significant using a two-sided paired t-test. Error bars indicates SD.

**Figure S4:**
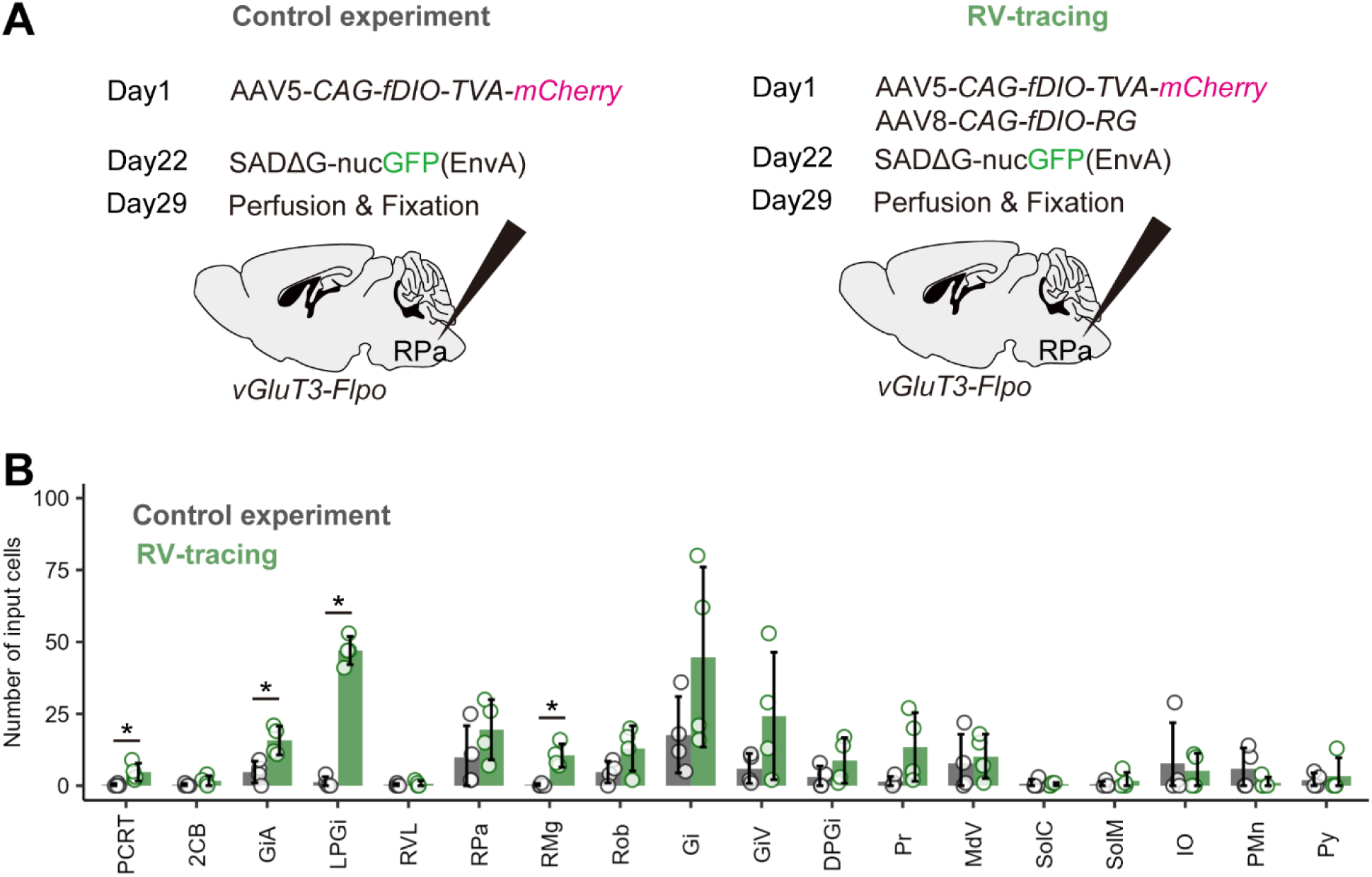
Control experiments for RV-mediated trans-synaptic tracing, related to Fig. 3. (A) Experimental procedure for RV-mediated trans-synaptic tracing and the control experiments. AAV8-*CAG-fDIO-RG* was omitted in the control experiment to assess the degree of RG-independent nonspecific infection^36^ of RV*ΔG-nGFP*+EnvA within and near the injection site. (B) The distribution of nGFP+ neurons found in both the control and RV-tracing experiments. *, *p* < 0.05 by two-sided exact rank sum test. N = 4 mice. For brain region abbreviations, see Table S1. Notably, we did not observe RV-nGFP labeling in the RG-omitted control group away from the injection site, such as in the hypothalamus, midbrain, or pons. In the medulla, we excluded areas with substantial nonspecific labeling from the analysis in Fig. 3, while utilizing the PCRT, GiA, LPGi, and RMg because trans-synaptically labeled neurons predominated in these areas.

**Figure S5:**
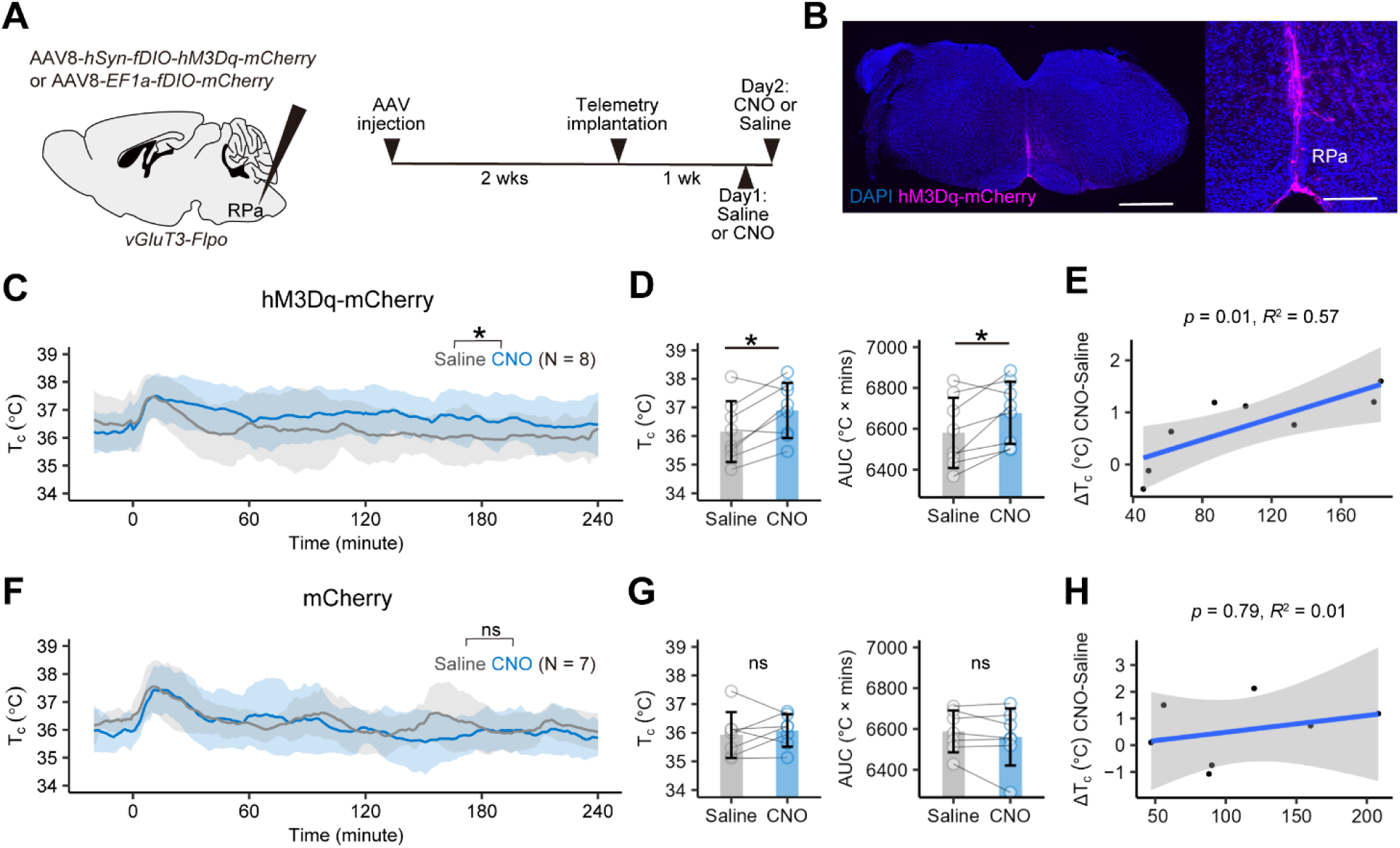
Chemogenetic activation of RPa-*vGluT3* neurons induces hyperthermia, related to Fig. 5. (A) Schematic of the virus injection and experimental timeline. AAV8*-hSyn-fDIO-hM3Dq-mCherry* or AAV8*-EF1a-fDIO-mCherry* was injected into the RPa of *vGluT3-Flpo* mice. (B) Representative image of hM3Dq expression in the RPa. The right panel shows an enlarged image. Scale bars, 1 mm (left) and 100 µm (right). (C, F) Group mean T_c_ traces in hM3Dq+ mice (C) or mCherry+ mice (F) following administration of saline (gray) or CNO (blue) at time 0. In panel C, repeated-measures two-way ANOVA showed significant solution (*p* < 0.01), time course (*p* < 0.01), and interaction (*p* < 0.01) effects. In panel F, only the significant time course effect (*p* < 0.01) is found. (D, G) T_c_ at 120 min (left) and AUC for T_c_ from 0 to 240 min (right) in hM3Dq+ mice (D) or mCherry+ (G) mice following saline or CNO injection. *, *p* < 0.05 by two-sided paired *t*-test. N = 8 for hM3Dq+ mice and N = 7 for mCherry+ mice. (E, H) Correlation between the number of hM3Dq-mCherry+ cells (E) or mCherry+ cells (H) and the change in T_c_ at 120 min following saline or CNO injection. The adjusted coefficient of determination (R^2^) is shown, with the *p*-value calculated using a *t*-test under the null hypothesis of no correlation. Error bars indicates SD.

**Figure S6:**
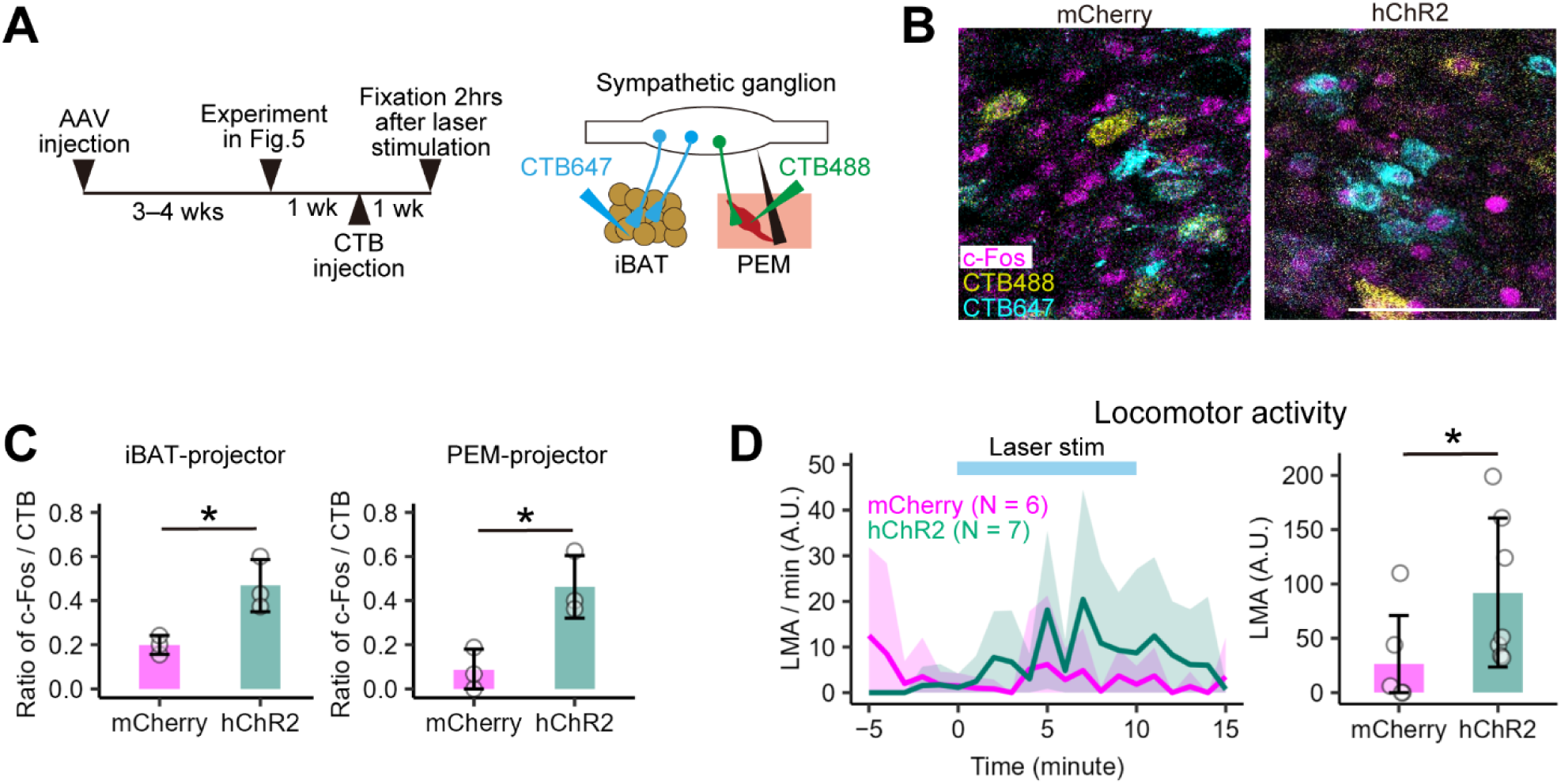
Additional information on activation of RPa-*vGluT3* neurons, related to Fig. 5. (A) Schematics for virus injections and the experimental timeline for c-Fos assay combined with retrograde labeling of the BAT-projecting and piloerector muscle-projecting sympathetic postganglionic neurons. This experiment is conducted following the data collection for Fig. 5. (B) Representative coronal section of the postganglionic neurons in the stellate ganglia. Magenta showing c-Fos immunostaining, while yellow and cyan denoting CTB-488 and CTB-647 labeling, respectively. Scale bars, 100 µm. (C) Fraction of c-Fos+ neurons among CTB-labeled neurons. *, *p* < 0.05 by two-sided unpaired student *t-*test. N = 3 for each group. (D) Count of locomotor activities following the laser stimulation. Right panel shows the total count of locomotor activities during the laser stimulation. *, *p* < 0.05 by Wilcoxon rank sum test. N = 6 for the mCherry group and N = 7 for the hChR2 group. Error bars, SD.

**Figure S7:**
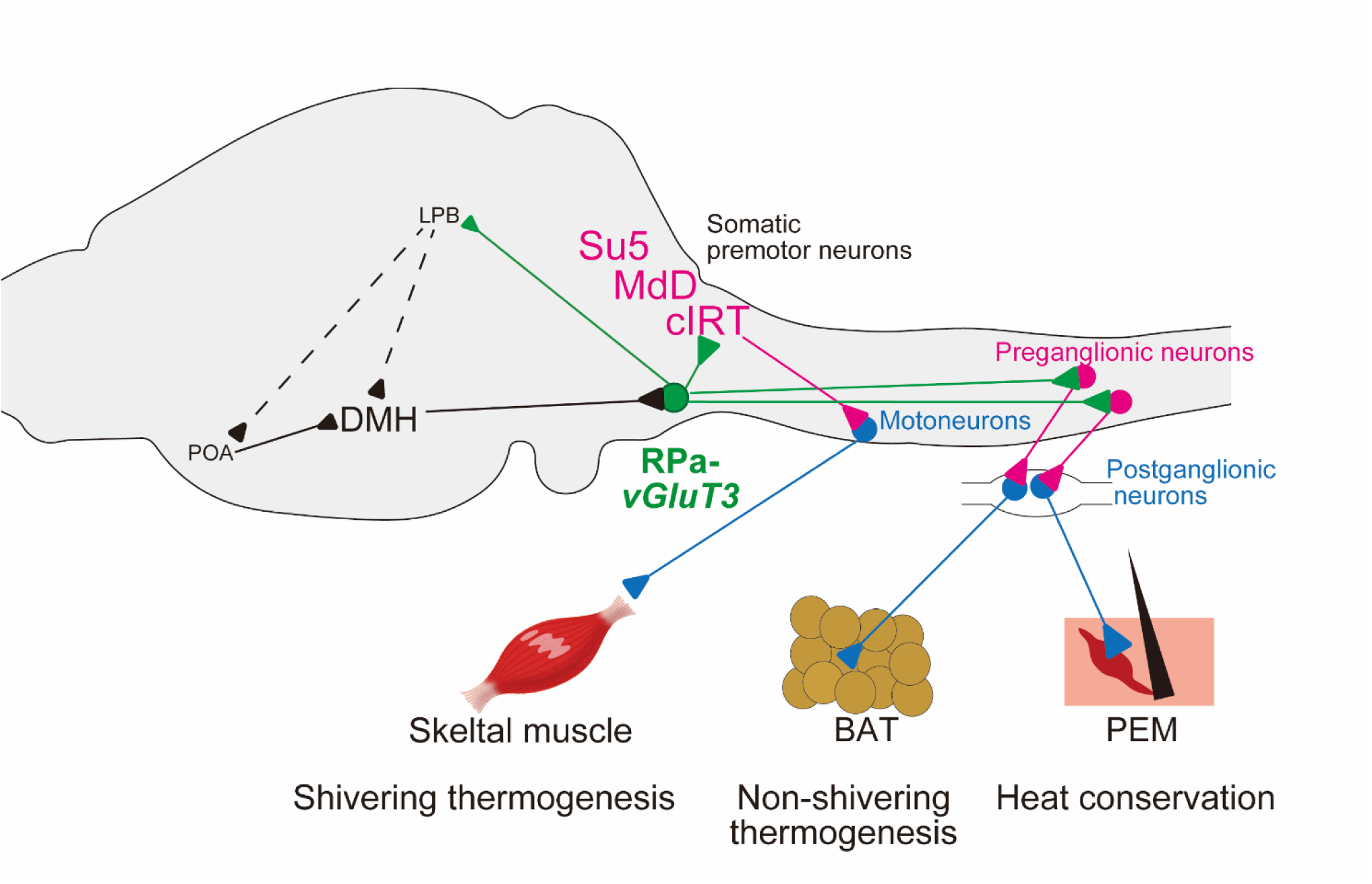
Proposed circuit model for thermogenesis regulated by RPa-*vGluT3* neurons. RPa-*vGluT3* neurons drive non-shivering thermogenesis in brown adipose tissue (BAT) and facilitate heat retention by engaging the piloerector muscle (PEM) via activation of the sympathetic nervous system. Concurrently, they recruit premotor neurons in the brainstem to initiate shivering thermogenesis through skeletal muscles. Although the RPa-*vGluT3* to LPB pathway, implicated in thermoregulatory behaviors, may contribute to feed-forward modulation of hypothalamic thermal centers, this specific circuit was not directly investigated in the present study.

